# Modelling human haematopoietic stem cell commitment *ex vivo* identifies IL-33 as a regulator of megakaryopoiesis

**DOI:** 10.64898/2026.09.01.748535

**Authors:** E.F. Calderbank-May, H.P. Bastos, L. Magnani, H. Foster, K. Sham, C. Wu, G. Mantica, C. Johnson, N. Mende, D. Hayler, S. Marra, J.S. Dahlin, C. Ghevaert, E. Laurenti

## Abstract

Commitment events to specific blood lineages arise from single hematopoietic stem cells (HSCs) and are influenced by stress, inflammation and disease. However, the understanding of how such events are regulated in human haematopoiesis is limited by the lack of tractable *in vitro* models. In this study, we introduce a novel Early Progenitor Differentiation (EPD) assay to study the initial lineage commitment of human 49f⁺ HSCs, in a system faithfully recapitulating cell states observed *in vivo*. Combining single cell -omics approaches and single-cell functional assays, we show that IL-33 acts directly on human 49f⁺ HSCs activating the MAPK pathway to enhance their commitment towards Megakaryocytic–Erythroid–Mast cell Progenitors and subsequently megakaryopoiesis. This occurs without affecting HSC self-renewal via accelerated establishment of chromatin programmes associated with Erythroid and Megakaryocyte and mast cells lineages. Our findings demonstrate the utility of the EPD model to identify molecular regulators of human HSC differentiation and uncover a new role of IL-33 in haematopoiesis.

## INTRODUCTION

Red blood cells and platelets are among the most abundant circulating cells in the human body, and their continuous replenishment depends on the tightly regulated activity of hematopoietic stem cells (HSCs) and multipotent progenitors (MPPs). These populations sustain lifelong haematopoiesis through a hierarchical yet dynamic process that balances self-renewal with lineage commitment.

Foundational insights into haematopoiesis have been historically derived from mouse models, but due to the clinical utility of human HSCs, the past decade has seen a growing emphasis on defining robust *in vitro* models of human haematopoiesis. First, there has been much progress in the development of *in vitro* models to expand HSCs to provide increased numbers of HSCs to patients undergoing transplantation or gene therapy (reviewed in (Bozhilov *et al*., 2023)). Second, directed differentiation protocols have been successfully established to generate red blood cells or platelets from cord blood-derived CD34⁺ cells, with the goal of producing clinically relevant transfusion products (Giarratana *et al*., 2011; Yang *et al*., 2016; Guan *et al*., 2017, 2020; Zhang *et al*., 2017; Fujiyama *et al*., 2020). Third, there has been a collective effort to develop culture conditions that sustain HSC differentiation to all blood lineages (Doulatov *et al*., 2010; Notta *et al*., 2016; Psaila *et al*., 2016; Belluschi *et al*., 2018; Tomei *et al*., 2025) as research tools to better understand the regulation of human haematopoiesis. However, most *in vitro* systems are tailored to HSC expansion or terminal differentiation, limiting their ability to resolve the earliest lineage choices downstream of HSCs. Although commitment toward granulocyte–monocyte progenitors (GMPs) has been extensively characterised, particularly in the context of emergency myelopoiesis (reviewed in (Swann, Olson and Passegué, 2024)) and the molecular regulation of erythroid (Ery) maturation at the BFU-E and CFU-E stages is well established (reviewed in (Schippel and Sharma, 2023)), the initial specification of Ery and megakaryocytic (Mk) fates from HSCs remains poorly defined. This disconnect highlights a key limitation in current human models, which do not adequately capture early fate decisions.

For many years, it has been postulated that all granulocytes (neutrophils, eosinophils (Eo), basophils (Ba) and mast cells (MC)) originate downstream of GMPs. However, recent evidence from single cell RNA-seq atlases of haematopoietic stem and progenitor cells (HSPCs) (Dahlin *et al*., 2018; Hay *et al*., 2018; Popescu *et al*., 2019) as well as single cell resolution functional assays (Drissen *et al*., 2019; Colin *et al*., 2026) has demonstrated that only neutrophils derive from GMPs. Eosinophils, basophils and mast cells instead are produced downstream of a progenitor which has lost lymphoid commitment capacity, but gives rise to mature Ery and Mk. Here we will refer to this progenitor using the terminology from Popescu et al., megakaryocyte–erythroid–mast cell progenitors (MEMPs) (Popescu *et al*., 2019). The MEMP compartment likely encompasses multiple previously described progenitor populations with overlapping lineage potential, including CD131⁺ common myeloid progenitors with basophil/MC/Ery/Mk output (Drissen *et al*., 2019), CD41⁺ megakaryocyte-biased progenitors (Miyawaki *et al*., 2017), and the F1–F3 megakaryocyte–erythroid progenitors (Notta *et al*., 2016). How the MEMP population and/or any of its subsets arise from HSCs, and how their production is dynamically regulated at steady state or during stress, remains unclear.

Here, we establish an early progenitor differentiation (EPD) *in vitro* model that recapitulates the earliest stages of human HSC differentiation into MEMPs. Integrating this system with *in vivo* approaches, we identify a novel role for the alarmin IL-33 in accelerating MEMP production, providing mechanistic insights into how inflammatory cues shape early lineage specification within the human hematopoietic hierarchy.

## RESULTS

### An *ex vivo* model for production of MEMPs from human HSCs

To model the earliest steps of differentiation from human HSCs, we cultured cord blood CD19^-^CD34^+^CD38^-^CD45RA^-^CD90^+^CD49f^+^ cells (49f^+^ HSCs), for 5 days in culture conditions that do not strongly drive differentiation (FLT-3, SCF, IL-3 and IL-6), in an assay we have termed Early Progenitor Differentiation (EPD). The time point of 5 days was selected as 49f^+^ HSCs will have completed an average of 2-3 divisions (Laurenti *et al*., 2015), and commitment to Ery and myeloid (My) lineages was observed in similar culture conditions from day 3 onwards (Johnson *et al*., 2024). To characterise the pool of HSPCs present in the culture at day 5 (hereafter termed d5-HSPCs), we performed xenotransplantation, single cell differentiation assays and single cell RNA sequencing (Figure 1A).

**Figure 1:**
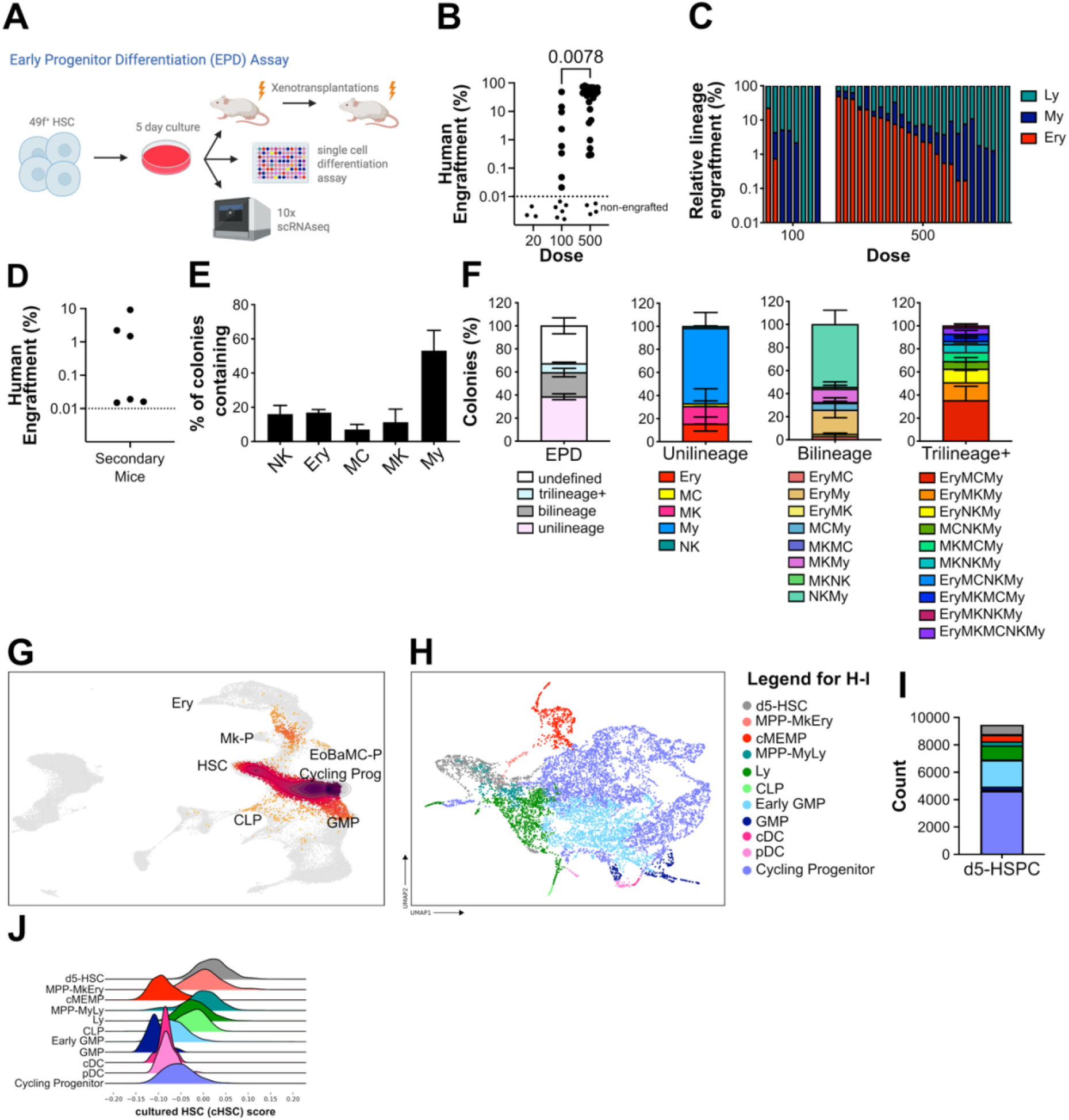
Development of an *in vitro* assay of Early Progenitor Differentiation (EPD). **(A)** Schematic of EPD assay. **(B-C)** Percentage of human engraftment (**B**) and relative lineage engraftment per mouse (**C**) in the injected femur of mice transplanted with d5 HSPC and then 20 weeks after transplantation (n= 4, 14 and 29 mice per dose). p values by Mann Whitney test. Dashed line: threshold of engraftment (%CD45^++^ + %GlyA^+^) ≥ 0.01 % and at least 30 cells recorded. Non-engrafted mice shown below dashed line. **(D)** Percentage human engraftment at 12 weeks post-secondary transplantation (n=6 mice). Mice were transplanted with CD34^+^ CD38^-^ cells isolated from primary transplanted mice shown in (B). Engrafted mice defined as above. **(E-F)** 49f^+^ HSCs were cultured 5 days in the EPD assay and then single d5-HSPCs were sorted into differentiation cultures. Percentage of colonies containing differentiated cells of the indicated lineages **(E)** or of the indicated type **(F)**. Data from n=3 independent CBs, n= 784 colonies. Mean ± SEM is shown. (**G**-**J)**: (**G**) Analysis of 10x genomics scRNAseq data from 49f^+^ cells cultured 5 days in the EPD assay (9,454 cells after QC). Projection on a published UMAP reference of a bone marrow dataset (Zeng *et al*., 2025). (**H**) UMAP with annotated cell clusters and (**I**) counts of each cluster. (**J**) Distribution of cultured HSC (cHSC) score (see methods) in each cluster of embedding shown in **(H)**.

First, we assessed if long-term self-renewing HSCs are still present at day 5 of EPD cultures. We performed a limiting dilution assay by intrafemoral injection of multiple doses of d5-HSPCs into sub-lethally irradiated immunocompromised NOD.Cg-Prkdc^SCID^Il2rg^tm1Wjl^/SzJ (NSG) mice. Flow cytometry analysis of bone marrow (BM) 20 weeks post-transplantation demonstrated the d5-HSPC population contains approximately 1:214 cells capable of long-term multilineage engraftment (Figure 1B-C, S1A). To evaluate self-renewal potential, CD34^+^CD38^-^ cells were flow-sorted from primary recipients and intrafemorally injected into secondary NSG mice. Successful human engraftment (Figure 1D, Supp Table 1) confirmed the presence of serially repopulating HSCs within EPD culture at day 5.

To reliably assess the differentiation potential of single cells within the EPD assay, we developed a single-cell functional assay capable of detecting all major haematopoietic lineages by modifying our previously published protocol (Belluschi et al., 2018), to include a step permissive for MC differentiation (see methods), enabling detection of Ery, Mk, MC, My and Lymphoid (Ly, natural killer (NK) cells) from single cells. We first validated this assay using single non-cultured 49f^+^ HSCs. After 3 weeks of culture, MCs were detectable by flow cytometry and cytospins (Figure S1B-E), alongside Ery, Mk, My and NK cells. When single d5-HSPCs were cultured in these conditions, they produced colonies containing a variety of combinations of Ery, Mk, MC, My and NK cells (Figure 1E-F and S1F). Notably, among the 57 colonies that produced MCs, 33 (57.9 %) also generated Ery cells and 10 (17.5 %) Mks (Figure S1G), suggesting that production of these lineages occurs via a tripotent Ery-Mk-MC progenitor in this model, similar to *in vivo* (Drissen *et al*., 2019).

To better characterise the progenitor cell types produced in the EPD model, 11,190 49f^+^ HSCs were cultured in the EPD assay and live d5-HSPCs were sorted for single cell RNAseq (10x Chromium technology) at day 5. Of the 10,668 cells sequenced, 9,454 cells passed QC. To estimate how the HSPCs produced in the EPD assay mapped to the continuum of haematopoietic cells observed *in vivo*, we projected these cells onto a high-resolution scRNA-seq reference dataset of BM haematopoiesis (Zeng *et al*., 2025). Projected d5-HSPCs spread from HSCs to Ery, Mk, Eo/Ba/MC, My and Ly progenitors (Figure 1G) indicating that EPD culture of 49f^+^ HSCs generates progenitor cells of all major haematopoietic branches.

For further analyses, we generated an independent UMAP embedding with 11 transcriptionally distinct cell populations, which we annotated using label transfer from a published reference dataset (Zeng *et al*., 2025) and subsequent manual curation (see Methods) (Figure 1H-I). Lineage-committed progenitor annotations were further validated using published lineage scores (Laurenti *et al*., 2013) (Figure S1H-I, Supp Table 2). State-of-the-art identification of the most immature HSCs in *in vivo* datasets relies on quantifying HSC dormancy signatures (Zhang *et al*., 2022). However once cultured all HSCs exit quiescence and enter cell cycle, which renders these signatures less reliable. We circumvented this problem by generating a transcriptional signature of cultured HSCs (cHSC signature) using published data comparing CD49f^+^ HSCs and CD34^+^ cultured for 3 days (Johnson *et al*., 2024) (see Methods). In line with the presence of serially transplantable HSCs at day 5 in EPD (Figure 1D), the cHSC signature was highest expressed in the day 5 HSCs (d5-HSC) cluster and then in the Ery/Mk-primed MPP (MPP^MkEry^) and My/Ly-primed (MPP^MyLy^) clusters, consistent with expected differentiation trajectories (Figure 1J). We observed that genes associated with Ery (*GATA1*, *KLF1*), Mk (HPGDS, *ITGA2*), and MC (*FCER1A*, *CPA3*) lineages were co-expressed within the same transcriptional cluster (Figure S1J). Based on this, we collectively refer to these cells as cultured Mk–Ery–MC progenitors (cMEMPs).

In summary, EPD culture conditions model the earliest differentiation steps of human HSCs with a 5-day window, recapitulating the transition from HSC to progenitor states observed *in vivo*, with demarked specification of Mk-Ery-MC progenitors.

### cMEMPs encompass Ery/Mk/MC progenitors and can be prospectively purified

To further investigate the transition from HSC to cMEMP, we compared differential gene expression between cMEMP, MPP^MkEry^ and d5-HSC populations. This analysis identified 7,653 genes significantly differentially expressed between d5-HSC and cMEMP (Supp Table 2). Gene set enrichment analysis (Laurenti *et al*., 2013; Velten *et al*., 2017; Hay *et al*., 2018; Mende *et al*., 2022) revealed enrichment for Ery and Mk lineage priming signatures in cMEMP compared to d5-HSC (Figure 2A, Supp Table 2). Genes associated with HSCs (*HLF*) and HSC/Ery priming (*CD52*, *MLLT3*) were downregulated with differentiation, whereas genes linked to Ery (*GATA1*/*KLF1*), Mk (*HPGDS*/*ITGA2B*) and MC (*FCER1A*/*CPA3*) lineages were significantly upregulated in cMEMP compared to MPP^MkEry^ and d5-HSC (Figure 2B). Given the early association between Ery, Mk and MC lineages (Dahlin *et al*., 2018; Hay *et al*., 2018; Drissen *et al*., 2019; Popescu *et al*., 2019), we next assessed the level of multilineage transcriptional priming of cMEMPs using lineage scores derived from published gene sets of highly purified HSPC subsets (Laurenti *et al*., 2013; Wu *et al*., 2022). 43% of single cMEMPs were transcriptionally primed for 2 or 3 of the Ery, Mk and MC lineage (Figure 2C, Supp Table 2). This indicates that cMEMPs likely constitute a mixture of uni-, bi-and tripotent progenitors of the Ery, Mk and/or MC lineages. To examine transcriptional pathways characterising cMEMPs, we performed gene set enrichment analysis comparing d5-HSC and cMEMP (Suppl. Table 2). Gene sets associated with signalling (including IL-1 and MAPK signalling), inflammation and cell cycle were significantly enriched in cMEMPs compared to d5-HSCs (Figure 2D).

**Figure 2:**
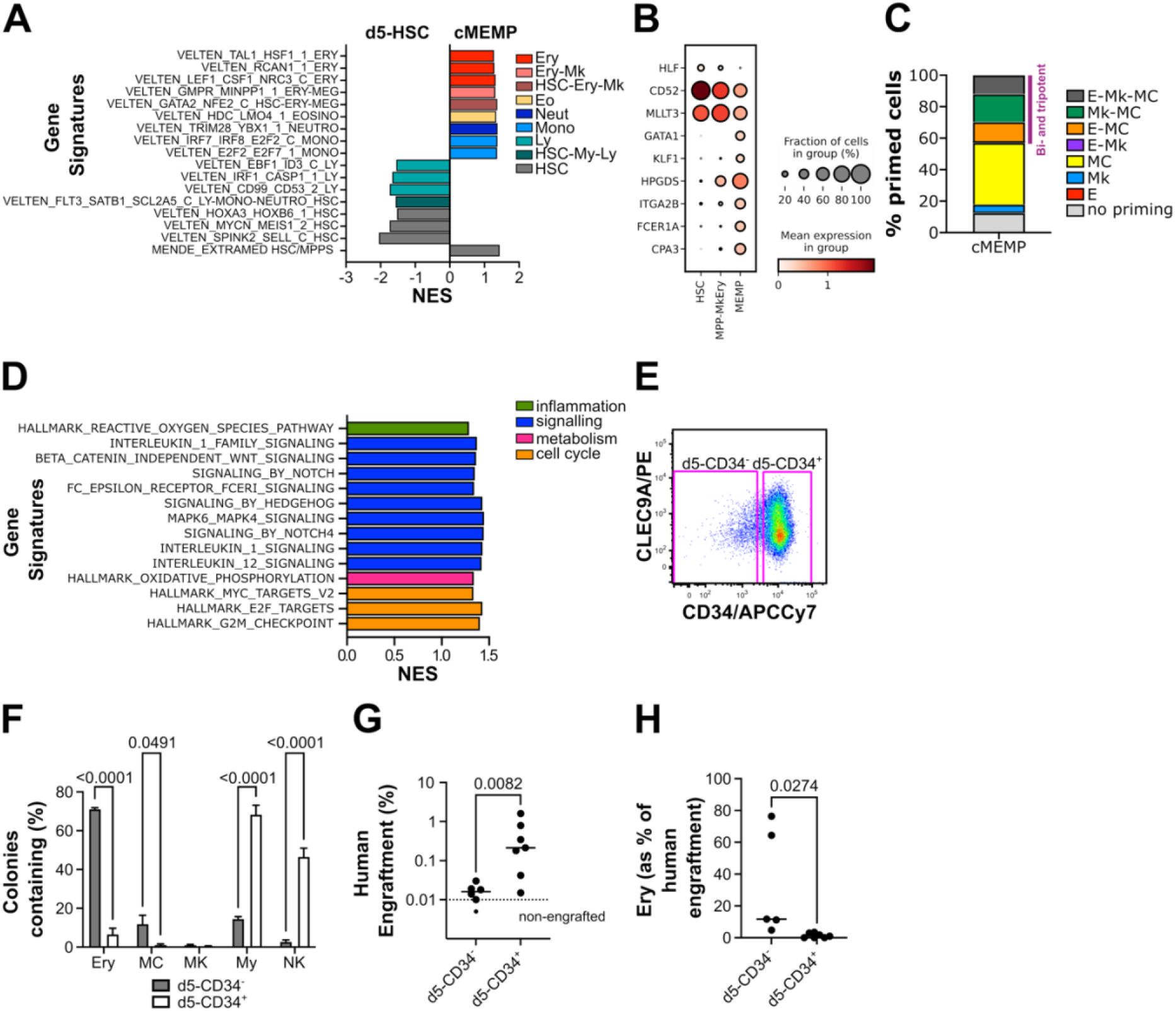
The EPD model generates prospectively purifiable cMEMPs. **(A**-**D)**: Analysis of 10x genomics scRNAseq of d5-HSPC from EPD culture. **(A)** Gene set enrichment analysis (GSEA) comparing d5-HSC and cMEMP of population-specific gene signatures from gene modules specific to the human HSC and progenitor cells indicated (gene sets from (Velten *et al*., 2017; Mende *et al*., 2022), selected gene sets with FDR<0.05 shown, NES: normalised enrichment score. **(B)** Expression values of selected marker genes in the HSC, MPP^MkEry^ and MEMP clusters. Circle colour shows mean expression values and circle size represents the proportion of cells expressing that gene per cluster. **(C)** Percentage of single cMEMPs displaying transcriptional priming towards the Ery, Mk and MC lineages (see methods). **(D)** GSEA comparing d5-HSC and cMEMP of selected Hallmark and Reactome genesets with FDR<0.05. (**E**) Representative flow cytometry plot showing gates used for d5-CD34^-^ and d5-CD34^+^. **(F)** Percentage of colonies containing differentiated cells of the indicated lineages, generated by d5-CD34^-^ and d5-CD34^+^ that were single cell sorted into differentiation cultures. Data from n=3 independent CBs, n= 464 and 687 colonies in d5-CD34^-^ and d5-CD34^+^ respectively. Mean ± SEM is shown, p<0.0001 by 2-way ANOVA with Sidak’s multiple comparisons (shown). (**G**-**H)**: Percentage of human engraftment (%CD45^++^ + %GlyA^+^) in the injected femur of mice transplanted with d5-CD34^-^ and d5-CD34^+^ cells at 2 weeks post transplantation (**G**) and percentage of erythroid cells (as percentage of human engraftment) (**H**). Data from n=6 mice for d5-CD34^-^ and n=7 mice for d5-CD34^+^. Dashed line: threshold of engraftment (%CD45^++^ + %GlyA^+^) ≥ 0.01 % and at least 30 cells recorded. Non-engrafted mice shown below dashed line, p values by Mann-Whitney test.

To identify a strategy to prospectively purify cMEMP cells *in vitro*, we used the index flow-sorting data from the d5-HSPC single cell differentiation assay (Figure 1E-F) and correlated cell surface marker expression at day 5 with the single cell colony outputs after an additional 3 weeks in culture (Supp Table 2). This analysis revealed significant differences in surface marker expression associated with colony type (Figure S2A). PCA loadings for PC2 indicated that cell surface expression of CD36, CD38, CD45RA and CD34 at day 5 were candidate markers for separating d5-HSPCs giving rise to colonies containing Ery, Mk and/or MC, compared to those giving rise to other mature cell types (Figure S2B). Based on the absolute levels of protein expression as well as the difference in expression between d5-HSPCs producing Ery, Mk and MC vs those producing other mature cell types (Figure S2C), we selected low cell surface expression of CD34 at day 5 as a potential prospective enrichment strategy for cMEMPs. To test this, we fractionated d5-HSPCs into CD34^-^ and CD34^+^ subpopulations (Figure 2E). Single d5-CD34^-^ or d5-CD34^+^ cells were sorted at day 5 of EPD culture into the differentiation assay described above and cultured for 3 weeks. Single d5 -CD34^-^ cells generated significantly more Ery and MC containing colonies than d5-CD34^+^ cells (Figure 2F, S2D-E). To assess their repopulation capacity, d5-CD34^-^ or d5-CD34^+^ were sorted and transplanted by intrafemoral injection into NSG mice. At 2 weeks post-transplantation, both populations engrafted, but d5-CD34^-^ engraftment levels were significantly lower than those of d5-CD34^+^ (Figure 2G and S2F, Supp Table 1) and contained a higher proportion of Ery cells (Figure 2H). In summary, actively proliferating progenitor cells with Ery, Mk and MC potential and limited engraftment capacity are generated by day 5 in the EPD model and can be prospectively isolated as CD34^-^ cells (henceforth referred to as phenotypic cMEMP).

### IL-33 increases cMEMP production via MAPK with no effect on proliferation or self-renewal

Enrichment of gene sets related to cytokine and inflammatory signalling in cMEMPs suggests that inflammatory cytokines may be regulating generation of this population and potentially their further differentiation. Several IL1 family genes were detected as differentially expressed in cMEMPs compared to d5-HSC or MPP^MkEry^ (Figure S3A), with *IL1RL1* standing out due to its strongest expression in cMEMP and MPP^MkEry^ (Figure 3A).

**Figure 3:**
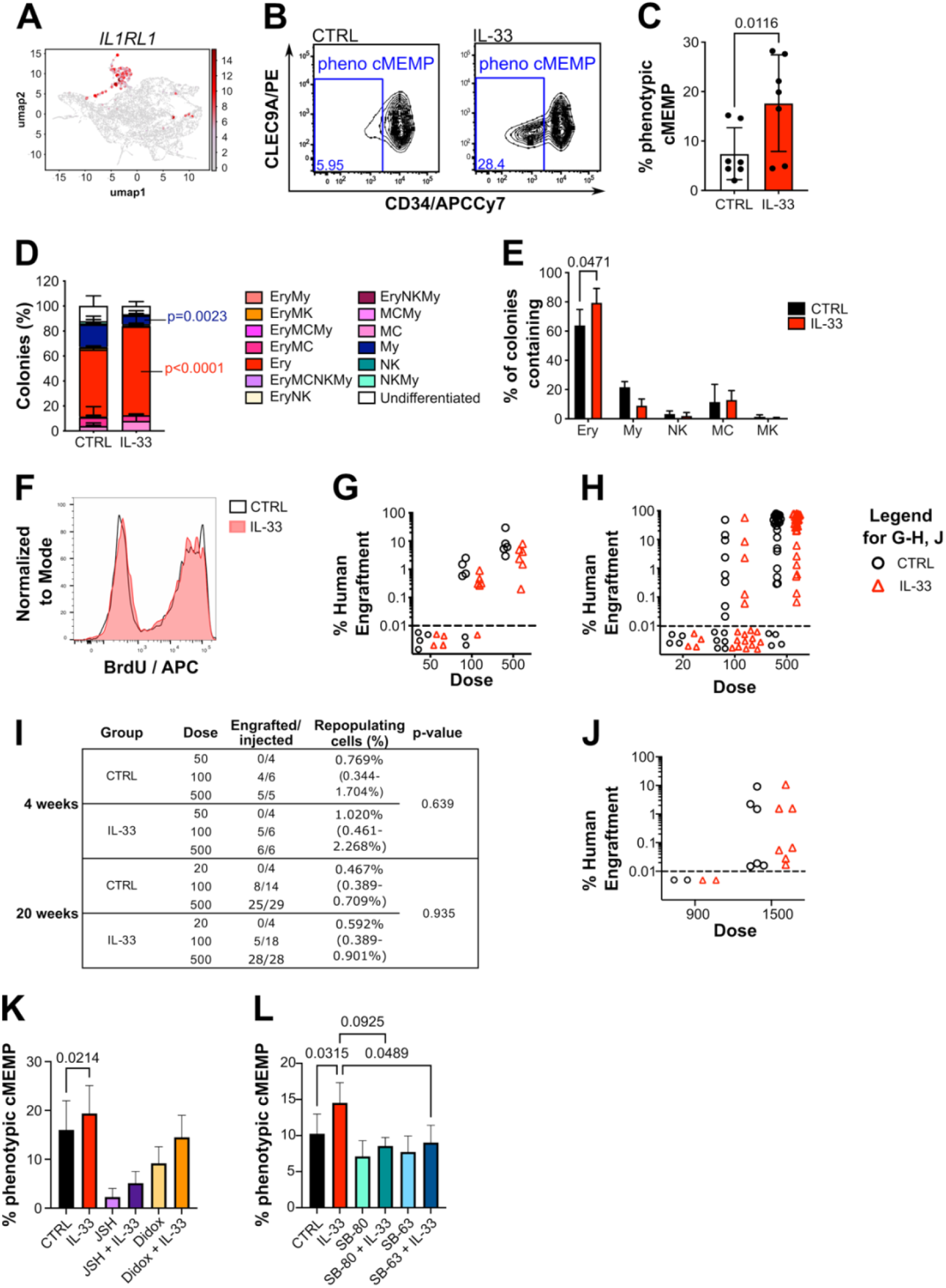
IL-33 influences cMEMPs production. (**A**) Expression of *IL1RL1* on d5-HSPCs shown on UMAP embedding from Fig.1H. (**B):** Representative flow plots of d5-HSPCs obtained from 49f^+^ HSC cultured 5 days in EPD ± IL-33. Blue box represents gate used for prospective purification of phenotypic cMEMPs. (**C**) percentage of phenotypic cMEMPs in presence or absence of IL-33 as assessed by flow cytometry, n= 7 independent CBs, p value by paired t-test. (**D-E):** percentage colonies of the indicated type (**D**) and percentage of colonies containing specified lineages (**E**) obtained from phenotypic cMEMP sorted at day 5 EPD ±IL-33 then cultured in same differentiation medium as in Figure S1D; n= 3 independent CBs, p=0.0261 by 2-way ANOVA with Sidak’s multiple comparisons (shown). (**F**) Representative BrdU staining of d5-HSPCs cultured ± IL-33. (**G-I)**: Percentage of human engraftment (%CD45^++^ + %GlyA^+^) in the injected femur of mice transplanted with d5-HPSC obtained from EPD cultures ± IL-33 at 4 weeks (**G**, n= 4, 6, 5 transplanted mice per dose respectively for CTRL and n= 4, 6, 6 transplanted mice per dose respectively for IL-33), and 20 weeks (**H**, n = 4, 14, 29 transplanted mice per dose respectively for CTRL and n= 4, 18, 28 transplanted mice per dose respectively for IL-33) after transplantation. Dashed line: threshold of engraftment (%CD45^++^ + %GlyA^+^) ≥ 0.01 % and at least 30 cells recorded. Non-engrafted mice shown below dashed line. CD45^++^ indicates cells positive for two distinct CD45 antibodies. **(I)** Frequency of long-term repopulating cells d5-HSPCs cultured ± IL-33, estimated using Extreme Limiting Dilution Analysis (ELDA) statistics. **(J)** Percentage of human engraftment (%CD45^++^ + %GlyA^+^) in the injected femur of mice transplanted with CD19^-^CD34^+^CD38^-^ cells isolated from primary transplanted mice (shown in **H**) at 12 weeks after transplantation (n = 3 independent experiments, 8 transplanted mice for CTRL and n= 9 transplanted mice for IL-33) Dashed line: threshold of engraftment (%CD45^++^ + %GlyA^+^) ≥ 0.01 % and at least 30 cells recorded. Non-engrafted mice shown below dashed line. CD45^++^ indicates cells positive for two distinct CD45 antibodies. **K-L**: Percentage phenotypic cMEMPs cells after 5-day culture of 49f^+^ HSC ± IL-33 with NFκB inhibitors, Didox (10 mM) and JSH-23 (25 mM) (**K**) and MAPK inhibitors, SB-203580 (SB-80, 5.3 mM) and SB-239063 (SB-63, 6.8 mM) (**L**); data shown ± SEM, n=3 independent CBs, p=0.0435 by mixed-effects analysis with sidak’s multiple comparisons (shown).

To determine whether IL-33 influences cMEMP production, we first added IL-33 (40 ng mL⁻¹) from day 0 of the EPD assay. IL-33 treatment significantly increased the proportion of phenotypic cMEMPs at day 5 compared with untreated controls (Figure 3B-C). To assess whether IL-33 alters lineage potential of cMEMPs, 49f⁺ HSCs were cultured in the EPD assay with or without IL-33, and at day 5 single phenotypic cMEMPs or d5-CD34⁺ cells (as in Figure 2E) were sorted into our multilineage differentiation assay in the absence of IL-33. In this assay, although the clonogenic efficiency (the proportion of single cells plated producing a colony) of single phenotypic cMEMPs was lower overall than that of d5-CD34⁺ cells, we observed comparable clonogenic efficiencies between control and IL-33 conditions (Figure S3B). IL-33 had no significant effect on colony composition derived from single d5-CD34⁺ cells (Figures S3C-D). Notably, IL-33 treated phenotypic cMEMP produced significantly more Ery-only and Ery-containing colonies and significantly fewer My-only colonies compared with untreated cMEMPs (Figures 3D-E).

Considering IL-33 accelerates the generation of more highly primed cMEMP from 49f⁺ HSCs, we next asked whether IL-33 affects proliferation or self-renewal. To assess proliferation, we performed a BrdU incorporation assay at day 5 and observed no difference between control and IL-33 treated cells (Figure 3F, S3E). Consistent with this, total cell numbers at day 5 were unchanged (Figure S3F). To evaluate self-renewal, d5-HSPCs from the EPD assay ± IL-33 were intrafemorally transplanted into sub-lethally irradiated NSG mice at multiple cell doses. *In vitro* IL-33 treatment had no effect on engraftment levels or lineage composition in primary recipients at 4 or 20 weeks post-transplantation, nor in secondary recipients at 12 weeks (Figure 3G-J, S3G-I, Supp Table 1). This indicates that IL-33 induction of cMEMP production occurs without altering HSC self-renewal.

IL-33 effects downstream of the ST2 receptor have been shown to be mediated by activation of either the IRAK1/MAPK and/or IRAK4/FRAF6/NFκB pathway in a cell type specific manner (Pusceddu, Dieplinger and Mueller, 2019). To functionally investigate by which signalling pathway IL-33 enhances differentiation towards cMEMP, MAPK inhibitors, SB 203580 and SB 239063, or NFκB inhibitors, JSH-23 and Didox, were added to the EPD assay in presence or absence of IL-33 and the percentage of phenotypic cMEMP cells was measured by flow cytometry. Both NFκB inhibitors decreased production of phenotypic cMEMPs in the absence of IL-33. However phenotypic cMEMPs production was increased by treatment with both JSH-23 and IL-33 or Didox and IL-33 (Figure 3K). This suggests that activation of NFκB contributes to cMEMP production *in vitro* but that the effect of IL-33 on cMEMP is NFκB independent. In contrast, treatment with both SB 203580 and IL-33 or SB 239063 and IL-33 returned the percentage of cMEMPs to that of the untreated condition therefore rescuing the effect of IL-33 (Figure 3L). This suggests that although NFκB plays a role in the production of cMEMP, IL-33 is acting via the MAPK pathway. Together these data demonstrate that IL-33 increases production of cMEMPs via the MAPK pathway, without changing proliferation or self-renewal of HSCs.

### IL-33 drives enhanced transcriptional lineage priming in cMEMPs

To further assess how IL-33 influences cMEMP production, we performed single-cell RNA sequencing (10x Chromium) on 49f⁺ HSCs cultured for 3 or 5 days in the EPD assay ± IL-33. At each time point, 20,000 live cells were sorted from control and IL-33 treated cultures. Using the same classification approach as in Figure 1H, we identified 10 transcriptionally distinct clusters (Figure 4A-B). Consistent with the proportional increase in phenotypic cMEMP observed above (Figure 3C), IL-33 treatment resulted in a higher proportion of transcriptionally defined cMEMP at both days 3 and 5 (Figure 4C).

**Figure 4:**
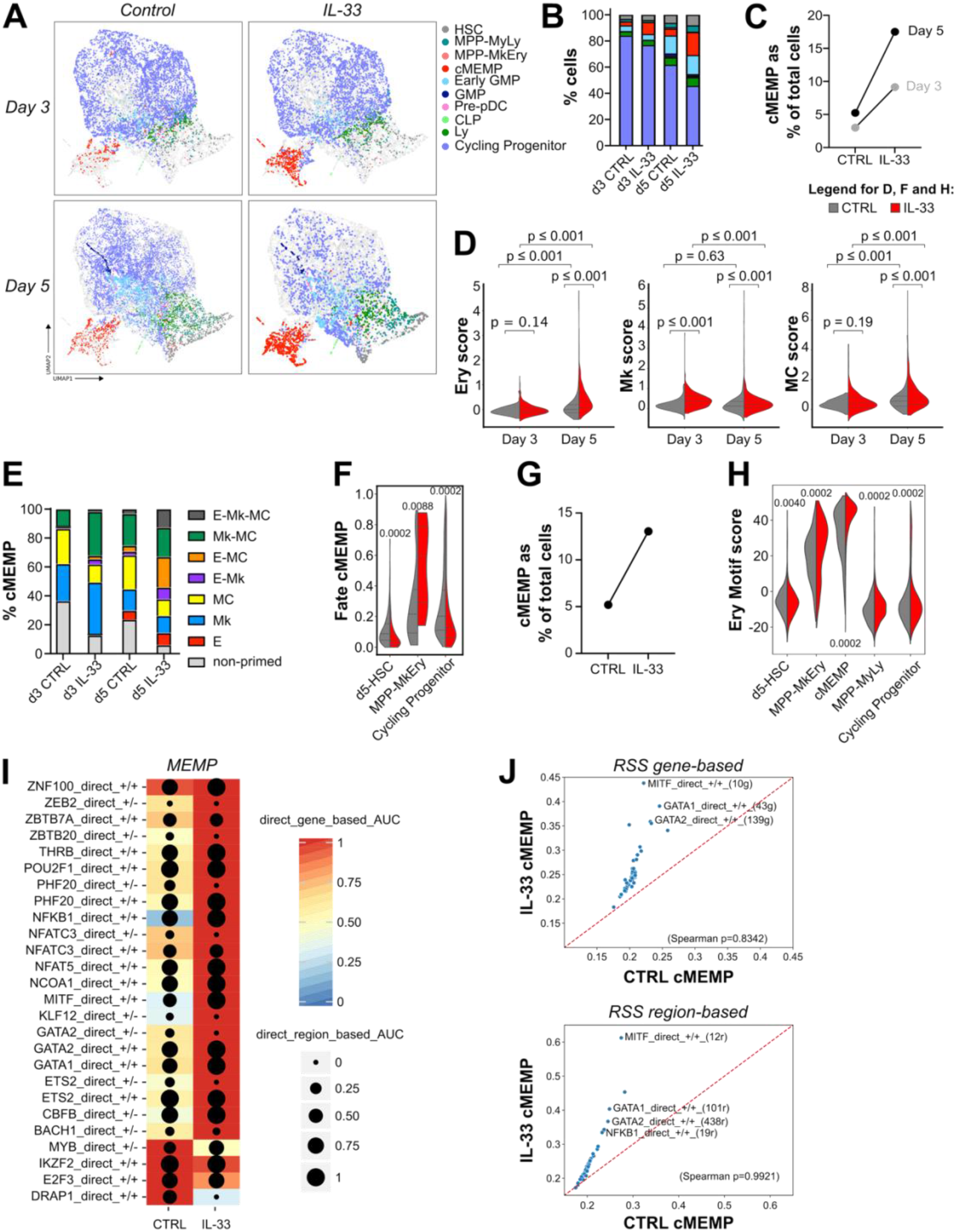
Amplification of cMEMP transcriptional networks by IL-33. (**A-F)**: Analysis of 10x genomics scRNAseq data from 49f^+^ HSCs cultured ± IL-33 in the EPD assay for 3 or 5 days. n= 4921, 4357, 8402 and 3048 cells in day 3 control, day 3 IL-33, day 5 control and day 5 IL-33 respectively (after QC). **(A)** UMAP embedding with annotated cell clusters. **(B)** Percentage of each annotated transcriptional cluster. (**C**) Percentage of transcriptionally defined cMEMP cells in the indicated conditions. (**D)** Ery, Mk and MC scores in cMEMPs in the indicated conditions. **(E)** percentage of single cMEMPs displaying transcriptional priming towards Ery, Mk and MC lineages (see methods for calculation). **(F)** Violin plots of FateID score in cells from the indicated clusters. (**G-J)**: Analysis of 10x multiome (combined scRNA-seq and scATAC-seq) of 49f^+^ HSCs cultured ± IL-33 in the EPD assay for 5 days. n= 17,703 and 15,928 cells in control and IL-33 conditions respectively (after QC). (**G**) Percentage of annotated cMEMP cells as annotated from the embedding shown in Fig S4E. **(H)** Violin plots of per-cell Stouffer scores calculated from the chromvar deviations computed on a set of motifs enriched in Erythroid-lineage cells (Takayama *et al*., 2021) for selected clusters. (**I**) Heatmap-dotplot for a selection of eRegulons contrasting CTRL vs IL-33 in cMEMPs. The heatmap colour represents regulon activity, the size of the dotplot represents regulon accessibility. **(J)** Regulon Specific Scores (RSS) scatter plots contrasting IL-33 treatment vs CTRL for the cMEMP cluster for gene-based (top) and region-based (bottom) eRegulons respectively. Red lines represent y=x. Selected eRegulons from the top 10 most group specific (as per RSS) highlighted with their respective labels. Statistics (Panels **D**, **F**, **H**): Internal dashed lines represent median and interquartile ranges. Group comparisons were evaluated using a two-tailed permutation of medians (10,000 iterations) test. Reported p-values are unadjusted for multiple corrections.

Pseudotime analysis of cMEMP suggested that at day 3, IL-33–treated cMEMPs likely progressed further along the differentiation trajectory than control cMEMPs (Figure S4A). To evaluate this further, we employed four complementary methods. First, we performed GSEA on a pre-ranked list of differentially expressed genes contrasting day 5 cMEMPs from IL-33 treated and control conditions, using published lineage-priming modules (Velten *et al*., 2017). IL-33–treated cMEMPs exhibited enrichment for Ery, Mk, eosinophil and myeloid priming compared with controls (Figure S4B, Supp Table 3). Second, we calculated “lineage scores” based on published lineage-specific gene sets (Laurenti *et al*., 2013; Wu *et al*., 2022) (see Methods). IL-33–treated cMEMPs showed significantly higher priming toward Mk and MC at day 3 and toward Ery, Mk, and MC at day 5 relative to controls (Figure 4D, Supp Table 3). Third, we assessed absence of priming as well as uni-, bi and tri-potent lineage priming towards Ery, Mk or MC as in Figure 2C. The proportion of primed cells was significantly higher in IL-33 treated cultures than in control cultures at both time points (Figure 4E, S4C).

To determine whether IL-33 increased multipotency within cMEMP, we compared uni-versus bi/tri-potent states (excluding non-primed cells). IL-33 treatment significantly increased the frequency of bi-and tri-potent cells at both days 3 and 5 (Figure 4E, S4D). Finally, we used the FateID algorithm to quantify cell lineage biases in progenitor populations (Herman, Sagar and Grün, 2018). This analysis revealed that IL-33 treatment increases the probability of MPP^MkEry^ cells to progress toward the cMEMP state (Figure 4F). Collectively, these data demonstrate that IL-33 accelerates cMEMP production and enhances transcriptional priming toward erythroid, megakaryocyte, and mast cell lineages.

### IL-33 amplifies the establishment of MEMP gene regulatory networks

Changes in chromatin accessibility precede expression of lineage-defining transcription factors and transcriptional signatures (Safi *et al*., 2022). To investigate how IL-33 influences chromatin accessibility during the HSC to cMEMP transition, we performed single-cell multiome profiling (Chromium Next GEM Single Cell Multiome ATAC + Gene Expression) on d5-HSPCs cultured in the EPD assay with or without IL-33. After quality control, cells were used to build an embedding based on the RNA modality. Clustering and annotation were performed as above (Figure S4E-F, Supp Table 4). Consistent with our previous dataset, IL-33 treatment increased the proportion of cMEMPs (Figure 4G), and these cMEMPs were more advanced along pseudotime than in the untreated condition (Figure S4G).

We next sought to uncover gene regulatory networks (GRNs) present in d5-HSPCs, using the SCENIC+ pipeline (Bravo González-Blas *et al*., 2023). This methodology uses the concordance between the expression of transcription factors (TFs) and that of their target genes with TF binding site accessibility to define an eRegulon for each TF (group of predicted target enhancers and predicted target genes). SCENIC+ identified expected TFs across specific HSPC subsets (*MEIS1* and *ERG* in HSCs, *IRF8* and *RUNX2* in DC-progenitors, *CEBPD* in GMP) and canonical lineage-defining regulons for Ery cells (*GATA1*, *BACH1*), Mks (*GATA1*, *ETS2*), and MCs (*MITF*) within the MEMP (Figure S4H). The enrichment of Ery related motifs in cMEMP was validated by a published Ery progenitor motif set (Takayama *et al*., 2021) being found significantly enriched in cMEMP, and to a less extent MPP^MkEry^ (Figure 4H). Notably, when comparing the untreated and IL-33 treated conditions, cMEMP-defining eRegulons exhibited increased chromatin accessibility upon IL-33 treatment (Figure 4I). Interestingly, IL-33 treated cMEMPs maintained largely the same set of eRegulons than control control cMEMPs but with higher specificities (Regulon Specific Scores, RSS) indicating increased activity of many lineage defining genes, based on both target gene expression and target gene accessibility (Figure 4J, Supp Table 4). These data demonstrate that IL-33 acts by amplifying the establishment of the cMEMP GRNs.

### IL-33 does not impact mast cell production downstream of unipotent mast cell progenitors

Given that IL-33 enhanced the production of Ery, Mk and MC primed cMEMPs, we next examined whether IL-33 equally promoted terminal erythroid, megakaryocyte, and mast cell differentiation downstream of cMEMPs. First, we focused on mast cell production. We used our newly established single cell differentiation assay supporting MC (described in Figure S1D) and plated single 49f^+^ HSC in presence or absence of IL-33. IL-33 significantly increased the percent of HSCs producing MC colonies (Figure 5A, S5A-B) but did not change the total number of MC produced per HSC (Figure S5C).

**Figure 5:**
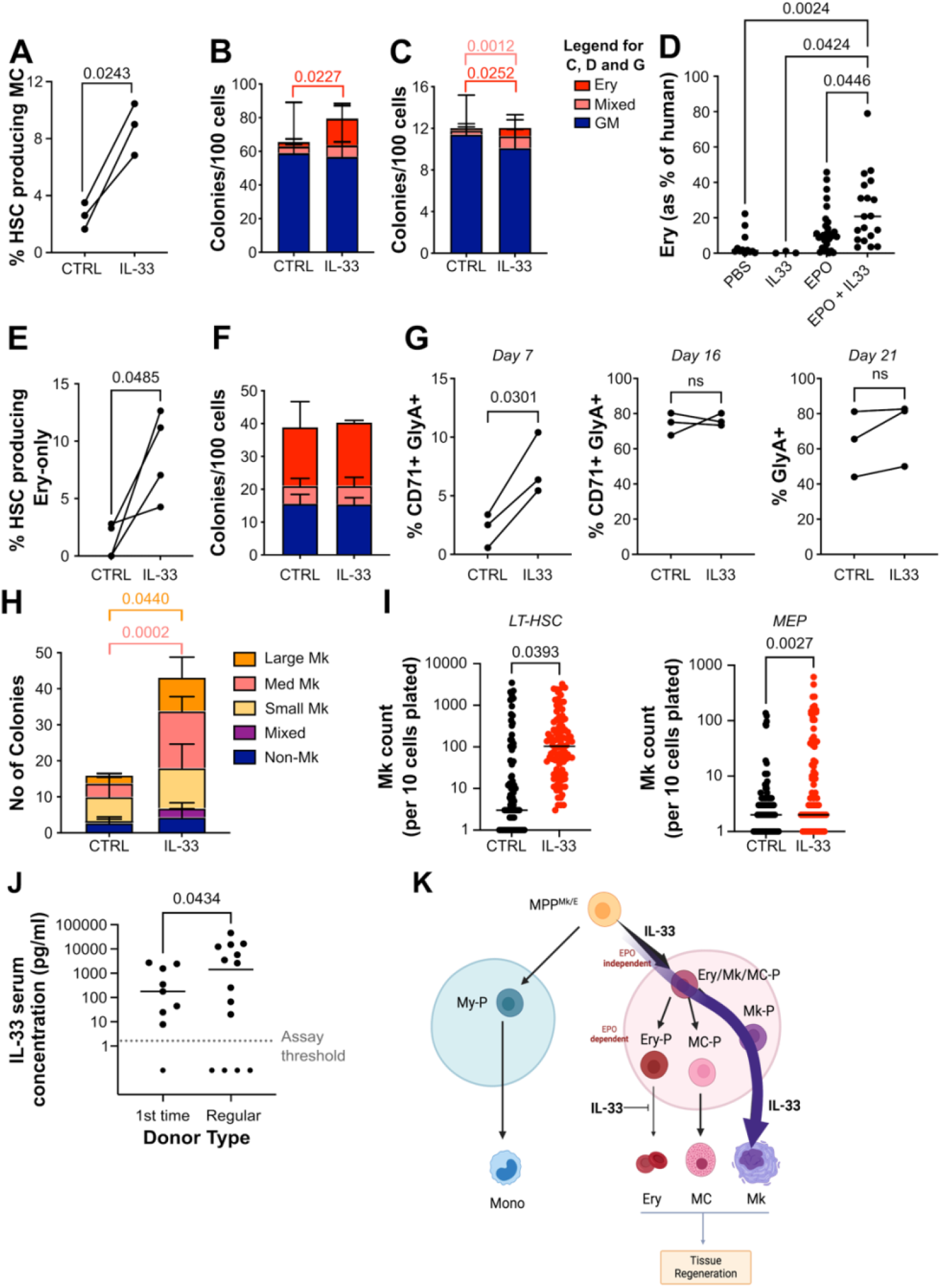
IL-33 affects both erythropoiesis and megakaryopoiesis. **(A)** Percentage of single cell 49f^+^ HSCs producing colonies containing MCs in differentiation medium as in Figure 1E (n=3 independent CBs, n=537 control and n=562 IL-33 colonies), p-value by paired t-test. (**B-C)**: Normalised number of colonies obtained in CFU assays initiated with either 49f^+^ HSC cultured for 5 days ±IL-33 in EPD assay (**B,** n=4 independent CBs) or day 0 49f^+^ HSCs (**C**, n=12 independent CBs). p values by paired t-test, mean ± SD shown. Colony types: Erythroid (Ery), granulocyte and monocyte (GM) or a combination of both (Mixed). (**D**) Ery engraftment as a percentage of human engraftment for NSG mice engrafted with CB CD34^+^ and then treated with PBS, IL-33, EPO or EPO and IL-33, n= 12, 3, 31 and 19 respectively. p=0.0013 by one-way ANOVA with Tukey’s multiple comparisons (shown). (**E**) Percentage of single cell 49f^+^ HSCs producing colonies containing only Ery in differentiation medium (n=4 independent CBs, n= 411 control and n= 445 IL-33 colonies), p-value by paired t-test. (**F**) Normalised number of colonies obtained in CFU assays initiated with CMP-MEP ± IL-33, n=3 independent CBs, mean ± SD shown. Colony types as above. (**G**) Red Blood Cell (RBC) assay: percentage of CD71^+^GlyA^+^ erythroblasts obtained at day 7 and 16 and GlyA^+^ erythroblasts at Day 21 after plating 500 49f^+^ HSCs at day 0 (n=3 independent CBs). p values by paired t-test. (**H**) Number of colonies generated by 750 49f^+^ HSCs plated into the MegaCult assay ± IL-33, n=4 control and n=6 IL-33 independent CBs, mean ± SD shown. p=0.0083 by 2-way ANOVA with Sidak’s multiple comparisons (shown), total counts of control vs IL-33 p=0.0143 by Mann-Whitney. (**I**) Number of Mk colonies obtained from 10 49f^+^ HSCs or MEPs plated into Mk differentiation medium (Psaila *et al*., 2016) ± IL-33 (n=1 CB, n= 96 colonies for each condition). p values by unpaired t-test. (**J**) Serum concentration of IL-33 from first time and regular platelet donors pre-donation (see methods), n= 9 and 14 donors respectively. Dotted line represents assay threshold (2 ρg mL⁻¹). p value by Mann-Whitney test. (**K**) Schematic of proposed effect of IL-33 in haematopoiesis.

### IL-33 promotes the early stages of erythropoiesis but inhibits terminal Ery differentiation

We next sought to assess at which stage of erythropoiesis IL-33 exerts its effects. First, 49f⁺ HSCs were cultured ± IL-33 for 5 days in the EPD assay. All d5-HSPCs were subsequently plated in a colony forming unit (CFU) assay for 2 weeks. IL-33 treatment resulted in an increased number of Ery and mixed (Ery and My) colonies (Figure 5B). A similar increase in Ery and mixed colony output was observed when day 0 49f⁺ and 49f⁻ HSCs were plated directly into a CFU assay with IL-33 (Figure 5C, S5D). Next, we checked the effects of *in vivo* IL-33 treatment. To this end, CB CD34^+^ cells were transplanted into NSG mice. 10 weeks post-transplantation IL-33 and/or EPO were injected intraperitoneally and BM composition was assessed by flow cytometry 2 weeks later. Whereas total human engraftment or B and My cell reconstitution were unchanged across conditions (Figure S5E-F), mice treated with EPO and IL-33 showed significantly elevated proportions of GlyA^+^ Ery cells than mice treated with EPO alone (Figure 5D, S5G, Supp Table 1).

To determine whether this increase in Ery output was driven by increased commitment to erythropoiesis or enhanced proliferation/maturation along that lineage, we cultured single 49f⁺ HSCs in a liquid-based differentiation assay that sustains simultaneous production of Ery and My cells (Belluschi *et al*., 2018). IL-33 production did not affect clonogenic efficiency (Figure S5H). The percentage of single 49f⁺ HSCs producing colonies containing Ery-only cells was significantly higher when the cultures were supplemented with IL-33 (Figure 5E, S5H-J). However, the number of Ery cells produced by each HSC was not affected by IL-33 (Figure S5K). These data suggest that increased Ery outputs upon IL-33 stimulation are most likely due to an effect on HSC commitment to cMEMPs rather than enhanced maturation downstream.

To test this hypothesis, we first plated d0 phenotypic MEMPs in a CFU assay in the presence or absence of IL-33. Consistent with IL-33 acting on HSCs rather than progenitors, we observed similar number of Ery colonies in control and IL-33 conditions (Figure 5F). Second, we cultured 49f⁺ HSCs ± IL-33 for 21 days in a liquid culture system enabling terminal erythroid differentiation (Lee et al., 2015), with flow cytometric analysis at day 7, 16 and 21. IL-33 treatment significantly increased the proportion of CD71^+^ GlyA^+^ erythroblasts in day 7 cultures (Figure 5G); but no differences in erythroblasts or mature cells were observed at later timepoints (Figures 5G). Interestingly, Swann et al. (Swann *et al*., 2020) reported that IL-33 inhibits terminal erythroid differentiation from committed erythroid progenitors, and here we independently confirmed this finding (Figure S5L). Together, these results demonstrate opposing roles for IL-33 during erythropoiesis: IL-33 enhances HSC commitment to erythroid progenitors while simultaneously inhibiting terminal erythroid maturation at later stages of erythropoiesis.

### IL-33 enhances megakaryopoiesis

Finally, we assessed how IL-33 perturbs megakaryopoiesis. Notably, 49f⁺ HSCs plated in semi-solid MegaCult medium with IL-33 generated a higher number of Mk colonies compared to untreated 49f⁺ HSCs, an effect primarily driven by a significant increase in medium and large sized colonies (Figure 5H). This suggests that IL-33 not only increases Mk commitment, but it may also enhance Mk maturation. To test this, we plated 10-cell mini-bulks in a liquid culture supporting Mk differentiation (Psaila *et al*., 2016). We observed that IL-33 increased total Mk output both from 49f⁺ HSCs (Figure 5I) and from downstream Megakaryocyte Erythroid progenitors (MEPs) (Figure 5I). These findings indicate that IL-33 enhances Mk production downstream of HSCs.

To investigate whether IL-33 may play a role in megakaryopoiesis in a human *in vivo* context, we thought to analyse IL-33 levels in individuals in which megakaryopoiesis is activated. We acquired serum from a cohort of platelet donors, including both regular donors, who have donated more than 25 times, and first-time donors. IL-33 ELISAs were performed on serum collected before the donation. Interestingly serum IL-33 levels were significantly higher in regular platelet donors than in first-time donors (Figure 5J). This indicates that IL-33 serum levels are elevated in individuals where emergency megakaryopoiesis is regularly elicited.

In summary, our study: i) establishes a novel model to examine the earliest divisions of human haematopoiesis, ii) identifies a common progenitor for erythroid, megakaryocyte, and mast cell lineages; and iii) reveals IL-33 as a previously unrecognised regulator of megakaryopoiesis, acting through coordinated induction of cMEMP from 49f⁺ HSCs (Figure 5K).

## DISCUSSION

Here we expand the experimental toolkit for human haematopoiesis by introducing a tractable model to study human HSC commitment to specific progenitor subtypes, which complements current efforts to expand HSCs *ex vivo* (Bozhilov *et al*., 2023; Sakurai *et al*., 2023) or sustain production of mature blood cell types (Doulatov *et al*., 2010; Notta *et al*., 2016; Belluschi *et al*., 2018; Drissen *et al*., 2019; Tomei *et al*., 2025). This method is scalable, amenable to screening using small molecule compounds, mRNA or gene editing and can also be used in disease contexts using primary patient samples. In this study, we demonstrate the utility of the EPD model to provide new mechanistic insights into the transition from HSC to MEMP and subsequent differentiation into Ery, Mk or MC and uncover IL-33 as a novel regulator of HSC early lineage commitment decisions.

### IL-33 induces megakaryopoiesis

Our study identifies IL-33 as a novel contributor of megakaryopoiesis. We observed that IL-33 exposure leads to a sustained increase in megakaryocyte output, accompanied by a reduction in erythroid cell production. Importantly, our data align with a previous report (Swann *et al*., 2020) to support a model in which reduced red blood cell production is not due to induction of a megakaryocytic bias at the level of cMEMP at the expense of entry into erythropoiesis. Rather, reduced red blood cell production results from a defect in Ery maturation, occurring post specification of unilineage Ery progenitors. Patients with thrombocytopenia have been found to either have lower levels of IL-33 in their bloodstream (Li *et al*., 2015) or harbour loss-of-function mutation in the *IL-33* gene (Lefrançais *et al*., 2026). Together with our observation of higher IL-33 levels in the serum of recurrent platelet donors, these data indicate a role for IL-33 signalling in promoting megakaryopoiesis and platelet biogenesis in humans at least under stress conditions.

### Coupling of stress megakaryopoiesis with emergency granulopoiesis via IL-33

Beyond their role of limiting blood loss upon vascular injury, platelets collaborate tightly with innate immune cells to ensure resolution of acute inflammation (Nicolai, Pekayvaz and Massberg, 2024). Therefore, coupling emergency granulopoiesis with stress megakaryopoiesis would maximise efficiency of haematopoietic recovery, contributing to keep platelet levels high during stress resolution and coordinating immune responses in injured, infected or inflamed tissues. We propose that the boost of cMEMP production induced by IL-33 from HSCs provides an example of how such coupling can be mechanistically achieved. This is in line with previous findings on the impact of IL-33 on haematopoiesis. First, knock-out (KO) of the IL-33 receptor *Il1rl1* in mice demonstrated direct regulation of mouse HSCs by IL-33 in transplantation settings (Capitano *et al*., 2020; Fu *et al*., 2025). Second, it is now clear that IL-33 is a key driver of regenerative responses in progenitor cells. IL-33 is released in the bone marrow by irradiation (Naef *et al*., 2025), 5-FU myeloablation (Fu *et al*., 2025) and helminth infection (Fagnan *et al*., 2026). Across all these settings, it generally drives progenitors into emergency granulopoiesis, with increased production of neutrophils, basophils and eosinophils (Kim *et al*., 2014; Fagnan *et al*., 2026). IL-33 treatment, like helminth infection, drives basophil and eosinophil differentiation by upregulating Lmo4 in basophil–eosinophil–mast cell progenitors within the erythroid-primed compartment (Fagnan *et al*., 2026), a population most likely downstream of the MEMPs described here. Neutropoiesis is also likely affected either directly (Kenswil *et al*., 2018; Fagnan *et al*., 2026) or indirectly through activation of type 2 innate lymphoid cells. Third, IL-33 shifts platelet function towards an inflammatory and granulocyte-recruiting phenotype as evidenced by proteomic studies (Gelon *et al*., 2026) and the fact that *IL-33* KO mice display defective neutrophil and eosinophil recruitment in intestinal (Chen *et al*., 2021) and lung (Takeda *et al*., 2016) inflammation models. Therefore IL-33 exerts complex actions across the spectrum of stem, progenitor, platelets and mature immune cells to promote recovery.

Finally, the mode of action of IL-33 reported here is in line with a lineage-instructive role. IL-33 enables more single 49f^+^ HSCs to acquire Ery, Mk and MC fates and accelerates the transition to MEMPs, without altering other cell fates (no loss of stem cell function, no decrease in commitment to other myeloid lineages such as monocytes, no overall cellular expansion). This inductive action seems fitting for a transient and efficient stress response resolution, likely enabling coordinated increased production of several cellular effectors of stress resolution. Future studies will have to address how chronic activation of such mode of action may contribute to pathogenesis in disease.

### Limitations

It is now well accepted that induction of megakaryopoiesis in mice can occur either via the classical route involving MPP^MkEry^ or a shorter route, directly branching of Mk-primed HSCs and bypassing MPP^MkEry^ (Sanjuan-Pla *et al*., 2013; Morcos *et al*., 2022; Poscablo *et al*., 2024; Meng *et al*., 2026). In this context, it is tempting to hypothesise that IL-33 induction of megakaryopoiesis via accelerated cMEMP production is akin to activation of the classical route instead of the by-pass one. Clonal tracking studies have documented the existence of Mk-primed HSCs in humans (Aksöz *et al*., 2024), but there are to date no markers to identify them and no evidence of a bypass megakaryocytic route in humans. We are therefore unable to formally prove this hypothesis.

In addition, cell surface expression of the IL-33 receptor IL1RL1 in human HSPCs cannot be robustly assessed due to the lack of reliable flow-cytometry antibodies. Interestingly, Il1rl1 is expressed on the surface of CD41^+^ Mk-biased mouse HSCs (Gelon *et al*., 2026). Future work will therefore have to address whether ILRL1 cell surface expression could enable purification of human Mk biased HSCs. Finally, while cMEMPs do resemble their *in vivo* counterparts and our study provides proof-of-principle that the EPD model can be used to identify new biology, like any *in vitro* models, it does not capture the complexity of the bone marrow microenvironment, and therefore may miss indirect effects driven by specific perturbations.

## MATERIAL AND METHODS

### 1. EXPERIMENTAL MODEL AND STUDY PARTICIPANT DETAILS

#### Umbilical cord blood collection

Umbilical cord blood (CB) samples were obtained from healthy donors with informed consent through the Cambridge Blood and Stem Cell Biobank (CBSB) or Anthony Nolan. All samples were collected according to procedures approved by the relevant Research Ethics Committees (07/MRE05/44, 18/EE/0199 and 24/EE/0116 research studies; IRAS Ref: 149581 for CBSB sourced material and 20/EE/0028 and 25/EM/0038 research studies; IRAS ref: 349351 for Nolan sourced material). CB samples were pooled irrespective of donor sex and processed as a single biological sample.

#### Peripheral blood

Peripheral blood (PB) cones were obtained from healthy donors with informed consent through CBSB. All samples were collected according to procedures approved by the relevant Research Ethics Committees (07/MRE05/44, 18/EE/0199 and 24/EE/0116 research studies; IRAS Ref: 149581).

First time or regular healthy male plateletpheresis donors were recruited and full consent given in accordance with East of England-Cambridge Central Research Ethics Committee (14/EE/0194).

#### In vivo studies

NOD.Cg-Prkdc^scid^Il2rg^tm1Wjl^/SzJ (NSG) mice were obtained from Charles River Laboratories or bred in-house. Experimental cohorts consisted of age-matched female mice (12–16 weeks old at the time of transplantation). All animals were maintained under Specific Pathogen-Free (SPF) conditions, and all procedures were performed in compliance with UK Home Office regulations. The study was conducted under the Animals (Scientific Procedures) Act 1986 Amendment Regulations 2012 following ethical approval by the University of Cambridge Animal Welfare and Ethical Review Body (AWERB).

## 2. METHOD DETAILS

### Isolation of CD34^+^ cells

Mononuclear cells (MNCs) were isolated by density gradient centrifugation using Lymphoprep (StemCell Technologies) or Pancoll (PanBiotech). Prior to separation, CB was diluted 1:1 with phosphate-buffered saline (PBS). Following centrifugation, the MNC layer was collected and subjected to red blood cell depletion using Red Blood Cell Lysis Buffer (BioLegend) for 15 minutes at 4 °C.

CD34⁺ hematopoietic stem and progenitor cells were positively selected from the MNC fraction using the CD34 MicroBead Kit (Miltenyi Biotec) and AutoMACS automated cell separation (Miltenyi Biotec) or manual MACS separation columns (Miltenyi Biotec), according to the manufacturer’s instructions. Purified CB CD34⁺ cells were cryopreserved and stored at −150 °C until use.

#### Flow cytometry and cell sorting

For cell sorting, cryopreserved CB CD34⁺ cells were thawed in pre-warmed IMDM (Life Technologies) supplemented with 0.1 mg mL⁻¹ DNase I (Lorne Laboratories) and 50% fetal bovine serum (FBS; PanBiotech). Cells were then resuspended in PBS containing 3% FBS and an antibody cocktail appropriate for the relevant experiment (Supp Table 5). Staining was performed for 20 min at room temperature (RT) in the dark, followed by washing with two volumes of PBS + 3% FBS.

Cells were sorted using a BD FACSAria Fusion or BD Influx cell sorter. For single-cell assays, sorting was performed in single-cell purity mode with index sorting enabled to permit retrospective linkage between cell surface phenotype and downstream colony outputs. Bulk sorts were performed in purity mode.

Previously defined hematopoietic populations were isolated, including 49f⁺ HSCs (CD19⁻CD34⁺CD38⁻CD90⁺CD49f⁺), 49f⁻ HSCs (CD19⁻CD34⁺CD38⁻CD90^-^CD49f^-^) and MEP (CD19^-^CD34^+^CD38^+^ CD45RA^-^CD123^-^), as well as additional populations described in the Results.

Flow cytometry analysis was performed on a BD LSRFortessa or BD LSRFortessa X-20. For analytical flow cytometry, cells were incubated with antibody panels in PBS + 3% FBS for 20 min at RT. All antibodies and complete staining panels are listed in Supp Table 5. Data analysed with FlowJo v9 or 10.

#### *In vitro* Early Progenitor Differentiation (EPD) Assay

For the EPD assay, Bulk-sorted 49f⁺ HSCs (1,500-2,000 cells) were plated in 96-well flat-bottom plates in a total volume of 200 µl StemSpan SFEM II (StemCell Technologies) supplemented with the cc100 cytokine cocktail (10 µl mL⁻¹; StemCell Technologies) and Penicillin–Streptomycin (1:50; Life Technologies), with or without recombinant IL-33 (40 ng mL⁻¹; Miltenyi Biotec). For inhibitor studies, cultures were additionally supplemented with SB203580 (5.3 µM), SB239063 (6.8 µM), JSH-23 (25 µM), or Didox (10 µM), with dimethyl sulfoxide (DMSO) used as a vehicle control. All inhibitors were obtained from Sigma-Aldrich.

#### Single cell Myeloid-Erythroid-Megakaryocte (MEM) assay

For the single cell MEM differentiation assay (as described in (Belluschi *et al*., 2018)), MS5 stromal cells were imported from Prof Katsuhiko Itoh at Kyoto University. One day prior to sorting, MS5 cells (passage 10–13) were seeded at 3,000 cells per well in flat-bottom 96-well plates in 100 µl Myelocult H5100 medium (StemCell Technologies) supplemented with 1% Penicillin–Streptomycin (Pen/Strep; Life Technologies) with or without recombinant IL-33 (40 ng mL⁻¹; Miltenyi Biotec).

On the day of sorting, culture medium was replaced with 100 µl per well of cytokine-rich MEM medium (StemPro medium with nutrient supplement, Life Technologies) supplemented with the following cytokines: SCF (100 ng mL⁻¹), FLT-3 (20 ng mL⁻¹), TPO (100 ng mL⁻¹), IL-6 (50 ng mL⁻¹), IL-3 (10 ng mL⁻¹), IL-11 (50 ng mL⁻¹), GM-CSF (20 ng mL⁻¹), IL-2 (10 ng mL⁻¹), IL-7 (20 ng mL⁻¹) (all Miltenyi Biotec), erythropoietin (EPO; 2 U mL⁻¹, Eprex, Janssen-Cilag), h-LDL (50 ng mL⁻¹; StemCell Technologies), 1% L-glutamine (Life Technologies), and 1% Pen/Strep. Single CB 49f⁺ HSCs were index-sorted directly onto the MS5 layer.

After 2 weeks of culture, colonies were harvested into 96-well U-bottom plates using a plate filter (Pall Laboratory) to remove MS5 cells. Colonies were stained for 20 min at room temperature in the dark with 50 µl of antibody mix (Supp Table 5), then washed with 100 µl PBS + 3% FBS. Colony type and size were quantified by high-throughput flow cytometry using the BD LSR II HTS Analyser. Erythroid colonies were identified as GlyA^+^, megakaryocytic colonies as CD41^+^, NK colonies as CD45^+^CD56^+^, monocyte colonies as CD45^+^ CD11b^+^ CD14^+^and granulocyte colonies as CD45^+^ CD11b^+^ CD15^+^. My defined as monocytes + granulocytes. Only colonies containing ≥30 singlet cells were included in downstream analysis.

#### Single cell Myeloid-Mast Cell-Erythroid-Megakaroycte (MCEM) assay

For the MCEM assay, single cells were sorted onto MS5 stromal cells (passage 8-10) as described above. After 7 days, 75 µl of MEM medium was removed and replaced with 75 µl MCEM medium, identical to MEM medium except for reduced SCF (5 ng mL⁻¹) and reduced EPO (1.5 U mL⁻¹). Colonies were harvested at 3 weeks post-sort and stained as above using the antibody panel listed in Supp Table 5. Gating strategy in Figure S1D. Cytospins were prepared to assist in morphological classification. Principal component analysis of colony output and index-sorting phenotypes was performed in GraphPad Prism (version 10). Values more that 5-times standard deviation were excluded (4 values total).

#### Colony-forming unit (CFU) assay

For the colony forming unit (CFU) assays, 500 49f⁺ HSCs (200 49f^+^ HSCs if cultured in EPD assay first) or 350 49f^-^ HSC or phenotypic MEMP (CD19^-^CD34^+^CD38^+^CD45RA^-^CD10^-^CD7^-^), were sorted into 500 µl PBS + 3% FBS in 1.5 mL microcentrifuge tubes. Cells were centrifuged (500 g, 5 min), resuspended in 50 µl PBS + 3% FBS, and mixed with 2.5 mL MethoCult Optimum (StemCell Technologies) supplemented with FLT-3 and IL-6 (10 ng mL⁻¹ each; Miltenyi Biotec). The mixture was plated at 1 mL per well into 6-well SmartDishes (StemCell Technologies). Plates were placed inside 24 cm² dishes surrounded by PBS and cultured for 2 weeks at 37 °C, 5% CO₂. Colony types (granulocyte-monocyte (GM), erythroid (Ery), or mixed (GM/Ery)) were scored manually at day 11 or using the StemVision instrument (StemCell Technologies) at day 14.

### *In vitro* erythropoiesis cultures

Erythroid differentiation cultures were performed using conditions adapted from Lee *et al*. (Lee *et al*., 2015). 500 49f⁺ HSCs cells were sorted and cultured for 7 days in Expansion medium (day 0–7), followed by sequential Differentiation medium I (day 7–12), Differentiation medium II (day 12–16) and Differentiation medium III (day 16-21). At the end of culture, cells were stained with the erythroid differentiation antibody panel (Supp Table 5) and analysed by flow cytometry.

To replicate the work of Swann (Swann *et al*., 2020), Mononuclear cells (MNCs) were isolated by density gradient centrifugation using Pancoll (PanBiotech) from PB cones. Prior to separation, the PB cones were draining into a 50 mL tube, diluted to 50 mL with PBS. Samples were centrifuged at 600g for 20 min, RT, acceleration 7, deceleration 0. Following centrifugation, the MNC layer was collected, washed and cells were cryopreserved and stored at −150 °C until use.

On the day of use, MNCs were thawed and CD34⁺ hematopoietic stem and progenitor cells were positively selected from the MNC fraction using the CD34 MicroBead Kit (Miltenyi Biotec) according to the manufacturer’s instructions. CD34^+^ cells were then stained with the antibodies for sorting (Supp Table 5). Live Lin^-^ CD34^+^ CD123^-^ CD71^+^ cells were sorted and plated 1,00 cells per well in StemSpan II (StemCell Technologies) + SCF (100ng mL⁻¹, Miltenyi Biotec) and EPO (2 units mL⁻¹, EPREX, Janssen) ± IL-33 (40 ng mL⁻¹, miltenyi). At day 8, cells were stained with (Supp Table 5) and analysed by HTS flow cytometry.

#### Megakaryocyte colony assays (Megacult-C)

Megakaryocyte colony-forming assays were performed using the Megacult™-C system with cytokine kit (StemCell Technologies), following manufacturer instructions. Briefly, 100 49f⁺ HSCs ± IL-33 (40 ng mL⁻¹) were mixed with 1 mL Megacult-C medium supplemented with 1% Pen/Strep, 40 µg mL⁻¹ h-LDL, and 0.6 mL Collagen Solution, and seeded in technical duplicates. Cultures were incubated for 12 days at 37 °C. Colonies were fixed in methanol:acetone (1:3) and stained for CD41 and nuclei according to the manufacturer’s protocol. Colonies were scored using a bright field microscope and identified as follows: small Mk colony: 3-20 CD41^+^ cells; intermediate Mk colony: 21-49 CD41^+^ cells; large Mk colony ≥ 50 CD41^+^ cells; mixed colony: ≥ 20 cells containing CD41^+^ and CD41^-^ (only nuclei staining) cells; non-Mk colonies ≥ 20 CD41^-^ cells. The average values of technical duplicates were used.

#### *In vitro* Megakaryocyte-Erythrocyte differentiation

Megakaryocytic cultures were carried out as in (Psaila *et al*., 2016). 10 MEP or 49f⁺ HSCs cells were sorted in 96 well plates and cultured for 9 or 13 days respectively. At the end of culture, cells were stained with the megakaryocytic differentiation antibody panel (Supp Table 5) and analysed by flow cytometry. Mk colonies were defined as CD41^+^ CD42b^+^.

#### BrdU Assay

To assess cell proliferation, 2,000 49f^+^ HSCs were cultured for 5 days in the EPD assay ± IL-33 (40ng mL⁻¹, Miltenyi). Cells were then fixed and stained for BrdU as manufacter’s protocol (BD Bioscience) and analysed by flow cytometry.

#### Xenograft experiments

All mice were sub-lethally irradiated (2.4 Gy) 24 hours before transplantation. Animals were anaesthetised with isoflurane and injected intrafemorally with the indicated doses of the specified cell populations, followed by subcutaneous administration of buprenorphine (0.1 mg kg⁻¹; Animalcare). At defined time points after transplantation, the injected femur was harvested and bone marrow (BM) was flushed. BM cells were stained using the antibody panels listed in Supp Table 5. To ensure robust detection of low-level human engraftment, two distinct anti-CD45 antibodies were used; cells were considered human if positive for both (CD45^++^). Mice were considered engrafted if (%CD45^++^+ %GlyA^+^) ≥ 0.01 % and comprised at least 30 events. Ery cells were identified as GlyA^+^, Ly cells as CD45^++^CD19^++^ and My cells as CD45^++^CD33^+^.

For limiting dilution analysis (LDA) experiments, 49f⁺ HSCs were cultured for 5 days in the EPD assay ± IL-33 (40 ng mL⁻¹, Miltenyi) before transplantation of d5-HSPCs. Mice analysed at 4, or 20 weeks post-transplantation received eight intraperitoneal injections of EPO (20 units per injection) during the 4 weeks preceding sacrifice.

To assess Ery potential following EPO and/or IL-33 treatment, NSG mice were transplanted intrafemorally with saturating doses of HSCs (43,000–69,000 CB CD34⁺ cells). After transplantation, mice received intraperitoneal injections every other day for 2 weeks of either PBS, EPO (20 units per injection; Eprex, Janssen), IL-33 (1 µg per injection; Miltenyi Biotec), or the combination of EPO and IL-33.

For secondary transplantation, CD34⁺ cells were purified from individual primary recipients using the CD34 MicroBead Kit (Miltenyi Biotec). CD19^-^CD34⁺CD38⁻ cells were subsequently isolated by flow cytometric sorting. Two doses of cells (5,000 or 900 cells) were transplanted into secondary recipients, which were analysed 12 weeks after secondary transplantation.

To assess the engraftment of d5-CD34^+/-^ cells, 5,000 d5-CD34^-^ or 20,000 d5-CD34^+^ cells were transplanted. Mice received four intraperitoneal injections of EPO (20 units per injection) and were sacrificed at 2 weeks post-transplantation.

### IL-33 ELISA

First time or regular healthy male plateletpheresis donors were recruited and full consent given. Platelet donations and sampling were carried out as previously described (Foster *et al*., 2020). IL-33 ELISA (Sigma-Aldrich) was carried out on pre-donation serum samples following manufacturer’s instruction. A threshold of 2 pg mL⁻¹ was used as the assay sensitivity.

#### scRNAseq and multiome assays

For single-cell RNA-sequencing experiments, 49f⁺ HSCs were cultured with or without IL-33 (40 ng mL⁻¹; Miltenyi Biotec) for 3 or 5 days. Following EPD culture, 20,000 live cells were sorted into 300 µl PBS + 3% FCS and kept on ice until library preparation. Cells were centrifuged and resuspended in 47 µl PBS + 0.04% BSA before processing with the Chromium™ Single Cell 3′ Library & Gel Bead Kit v3 (10x Genomics), according to the manufacturer’s instructions. Libraries were sequenced on one lane of a NovaSeq 6,000 instrument at the CRUK Cambridge Institute Genomics Core.

For single-cell multiome (ATAC + gene expression) profiling, 49f⁺ HSCs were cultured with or without IL-33 (40 ng mL⁻¹) for 5 days in the EPD assay. A total of 100,000 live cells were sorted into 300 µl PBS + 3% FCS and maintained on ice prior to library preparation. Samples were processed using the Chromium Next GEM Single Cell Multiome ATAC and Gene Expression workflow (10x Genomics), following the manufacturer’s protocol. Libraries were sequenced in one lane (per modality) on a NovaSeq 6,000 at the CRUK Cambridge Institute Genomics Core.

#### Reads demultiplexing, aligning and UMI quantification

The resulting sequencing outputs provided by the sequencing centre as FASTQ files were subsequently aligned to the GRCh37 (hg19) reference (refdata-cellranger-hg19-3.0.0 - Ensembl 87) and quantified through the use of *cellranger count* from the 10x Genomics Cell Ranger pipeline (Zheng *et al*., 2017) for Dataset 1 and Dataset 2 (versions 3.1.0 and 5.0.0 respectively). For the multiome (GEX+ATAC) experiment (Dataset 3) the output was aligned to the GRCh38 (hg38) reference (refdata-cellranger-arc-GRCh38-2020-A-2.0.0) and quantified using *cellranger-arc count* from the Cell Ranger ARC pipeline (version 2.0.1).

#### Filtering, normalization and dimensionality reduction

For the libraries processed by the Cellranger pipelines the Python package Scrublet (Wolock, Lopez and Klein, 2019) was subsequently used to estimate and remove putative multiplets. Further filtering and downstream processing was applied using the SCANPY toolkit (Wolf, Angerer and Theis, 2018). For the Gene EXpression (GEX) libraries, barcodes (cells) with less than 500 genes or which had their total gene expression over 10% or more of mitochondrial origin were also excluded. Only genes expressed in at least by three barcodes (cells) were retained. UMI counts were then log-normalised and a set of highly variable genes determined. Each dataset (Dataset 1 and Dataset 2) counts matrix was regressed for number of counts and fraction of mitochondrial counts and scaled before computing the Principal Components. The diffusion map (Haghverdi, Buettner and Theis, 2015) was calculated and its component representation used to generate a neighbors graph. Finally, UMAP (McInnes *et al*., 2018) embedding representations were also computed for each dataset.

*Dataset 1* (d5-HSPC RNA): from the 10,668 cells estimated by the Cell Ranger pipeline, 9454 cells passed quality control.

*Dataset 2* (d3/5-HSPC ± IL-33 RNA): Of the 5,306 (d3-CTRL), 5,006 (d3-IL-33), 9,778 (d5-CTRL), and 3,216 (d5-IL-33) cells estimated by the Cell Ranger pipeline, 4,921, 4,357, 8,402 and 3,048 cells, respectively, passed quality control.

*Dataset 3* (d5-HSPC ± IL-33 RNA + ATAC): After quality control, 17,703 control cells and 15,928 IL-33–treated cells were used to build an UMAP embedding based on the RNA modality.

#### Annotation transfer, pseudotime and clustering

A single-cell transcriptional atlas of human haematopoiesis (Zeng *et al*., 2025) was used to annotate our single-cell RNAseq datasets. Following the instructions at https://github.com/andygxzeng/BoneMarrowMap - which wrap around the R package of Symphony (Kang *et al*., 2021)- a reference mapping for each dataset was performed. Alongside it predicted pseudotime were also generated from the pipeline and assigned to our datasets. The transferred annotation labels were included into each dataset which was then subjected to high-resolution (Dataset 1/2: 25.0; Dataset 3: 15.0) clustering using the Leiden algorithm (Traag, Waltman and van Eck, 2019). The most frequent (with ties manually resolved) annotation in each of the resulting clusters was then adopted. Annotation label strings were adjusted to agglutinate closely related groups into a granularity of interest (Dataset 1: 11 clusters, Dataset 2: 12 clusters, Dataset 3: 9 clusters).

#### Donor demultiplexing

For RNAseq only datasets Dataset 1 and Dataset 2 donors were demultiplexed using cellSNP-lite (Huang and Huang, 2021) and vireo (Huang, McCarthy and Stegle, 2019) using the recommend settings and respective number of of donors for each dataset.

#### Differential expression & GSEA

Differential expression testing was performed with R NEBULA package (He *et al*., 2021). The Benjamini-Hochberg procedure (as implemented in Python *statsmodels* module) was applied to control the false discovery rate (alpha = 0.05). Rank files to be used with GSEA were also created for each computed contrast by taking the product of the sign of log-fold change and negative of log10 of the p-value.Gene set enrichment analysis (GSEA) was performed with the GSEA software (v4.3). Enrichment was tested for C2 curated gene sets and hallmark gene sets, published population-specific signatures (Laurenti *et al*., 2013; Hay *et al*., 2018) and lineage priming modules (Velten *et al*., 2017).

#### cultured HSC score

Samples of cultured mobilized peripheral blood comprising of CD34^+^ and LT-HSC cells were retrieved from https://www.ncbi.nlm.nih.gov/geo/query/acc.cgi?acc=GSE213372. The resulting dataset was filtered, normalised and differential expression was tested using DESeq2 (Love, Huber and Anders, 2014) for the cell type contrast using single-cell recommendations (https://bioconductor.org/packages/devel/bioc/vignettes/DESeq2/inst/doc/DESeq2.html#recommendations-for-single-cell-analysis) and accounting for the sequencing run and number of genes as covariates. Through examination of the differential expression results a log-fold change of 1.5 at a statistical significance of 0.01 were used as optimal cutoffs and yielded a gene set characteristic for cultured LT-HSC.

#### Lineage priming scores

Lineage priming scores and methodology were from (Mende *et al*., 2022). For the Ery, Mk and MC score thresholds, the 95^th^ percentile score was taken for d5-HSC and used as a negative threshold.

#### FateID

FateID (Herman, Sagar and Grün, 2018) was applied to Dataset 2 to compute fate bias probabilities for the cell types MEMP, GMP and Lymphoid. The resulting fate probabilities were visualized as split (treatment vs control) violins and a two-tailed median permutation test (10,000 iterations) performed to assert for significant differences.

#### pycistopic + SCENIC+

The multiome dataset (Dataset 3) was processed using the SCENIC+ pipeline (Bravo González-Blas *et al*., 2023) (version 1.0a2). Initially, the RNA-modality of Dataset 3 was processed similarly to the previous datasets as described above. The the ATAC-modality pycisTopic (included with SCENIC+) was used for QC, topic generation via Mallet (McCallum and Kachites, 2002) (version 2.0), imputing region accessibilities and differentially accessible regions. The processed modalities were then combined into a scenic plus object and automatically processed downstream by the pipeline.

#### Regulon Specificity Scores (RSS)

The RSS resulting from the SCENIC+ were inspected through the package native visualisation methods. In addition, scatter plots of RSS for the cMEMP cluster cells contrasting IL-33 treatment versus control were produced separately for the gene-based and region-based eRegulons. Spearman correlation of the contrasts was calculated.

#### Deviation scores (chromVAR) analysis

For comparing motif accessibility within Dataset 3 cells (chromVAR) deviation scores (Schep *et al*., 2017) were computed. The package scPrinter (Hu *et al*., 2025) was used to that effect to take advantage of the GPU implementation of the chromVAR method. As for motifs of interest the motif collection in the Takayama et al, 2021 study was extracted. Using Biopython tools (https://biopython.org/) the consensus sequences from that study were converted into a JASPAR file with a position-weighted matrices per motif that was used by scPrinter. The top scoring motifs (alpha = 0.05) for the Erythroid signature in the Takayama study were used to search for and contrast erythroid motif signature regions in Dataset 3. For that purpose, the deviations scores were converted into Stouffer scores (Stouffer *et al*., 1949) and contrasts were assessed for significance using a two-tailed permutation of medians test (10,000 iterations).

## 3. QUANTIFICATION AND STATISTICAL ANALYSIS

All of the statistical details can be found in the figure legend and methods. Significance was defined as p<0.05.

## ACKNOWLEDGMENTS

We would like to express our enormous gratitude for the generous donation of human tissue by the CB and PB donors. We would also like to thank the Cambridge NIHR BRC Cell Phenotyping Hub for their flow cytometry services and advice, the CRUK Cambridge Institute genomics centre for the RNA-seq library preparation sequencing and the staff of AMB for their support. This research was also supported by the CIMR Flow Cytometry Core Facility. We wish to thank Reiner Shulte and Gabriela Grondys-Kotarba for their advice and support in cell sorting. We thank Joanna Baxter, the team of the Cambridge Stem Cell Biobank for their support in sample management and acknowledge Anthony Nolan charity for provision of cord blood. We would like to thank Dr Juan Li and Dr Matthew Williams for general advice on the study.

## FUNDING

Wellcome – Royal Society Sir Henry Dale Fellowship 107630/Z/15/Z (EL)

Wellcome Discovery Award 309075/Z/24/Z (EL)

Kay Kendall Leukaemia Fund - KKL1325 (EL)

The Alborada Trust (EL)

Cancer Research UK Cambridge Cancer Centre PhD fellowship CTRQQR-2021 (GM)

Deutsche Forschungsgemeinschaft (DFG) Research Fellowship ME 5209/1-1 (NM)

This research was funded in whole, or in part, by the Wellcome Trust [203151/Z/16/Z, 203151/A/16/Z, 107630/Z/15/Z, 309075/Z/24/Z] and the UKRI Medical Research Council [MC_PC_17230]. **For the purpose of open access, the author has applied a CC BY public copyright licence to any Author Accepted Manuscript version arising from this submission.**

## CONTRIBUTIONS

Conceptualization: EL, EFCM

Methodology: EL, EFCM, JSD, CW, HPB

Investigation: EFCM, LM, HF, CW, GM, CJ, NM, DH, SM, JSD

Visualization: HPB, EFCM, CW

Formal analysis: HPB, EFCM, KS, CW

Data curation: HPB, EFCM

Validation: EL, EFCM Resources: CG

Writing—original draft: EFCM, EL

Writing—review & editing: EL, EFCM, HPB

Supervision: EL, EFCM

Project management: EL, EFCM, CG, JSD

Funding acquisition: EL

