## Supplementary Material for "Modelling human haematopoietic stem cell commitment *ex vivo* identifies IL-33 as a regulator of megakaryopoiesis"

### Supplementary Materials

**Supplementary Figures 1-5**
**Supplementary Tables 1-5**

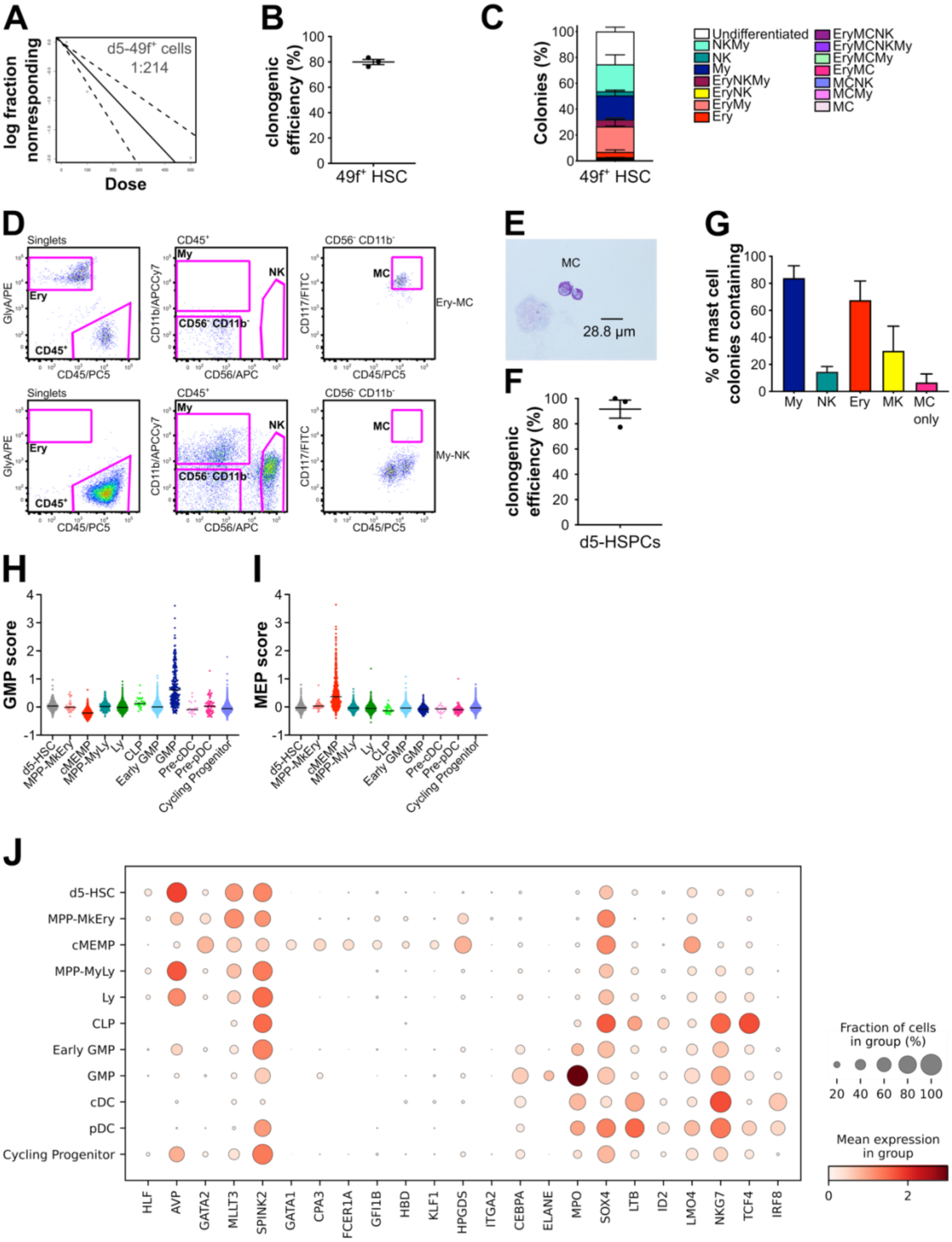


**Suppl. Figure 1: Characterisation of the Early Progenitor Differentiation Assay.** (**A**) Estimation of frequency of long-term repopulating cells within EPD d5-HSPCs. Analysed performed by ELDA with engraftment data at 20 weeks post-transplantation. (**B-E)**: day 0 49f+ HSCs were sorted into differentiation cultures enabling mast cell differentiation. (**B**) Clonogenic efficiency, (**C**) percentage of each type of colony produced, (**D**) representative flow cytometry plots of Ery-MC and My-NK colonies and (**E**) representative cytospin of mast cells generated in these cultures. Data from n=3 independent CBs, n= 537 colonies. Mean ± SEM is shown. (**F**-**G)**: d5-HSPCs generated in the EPD assay were single cell sorted and cultured in same conditions as above. (**F**) Clonogenic efficiency and (**G**) percentage of mast cell colonies containing different lineages. Data from n=3 independent CBs, n= 784 colonies. Mean ± SEM is shown. **(H**-**I)**: violin plots of (**H**) GMP and (**I**) MEP scores calculated for each cell in the indicated clusters. Gene sets from (Mende *et al.*, 2022). Expression of selected marker genes. **(J)** Circle colour shows mean expression values and circle size represents the proportion of cells expressing cells per cluster of 10x genomics scRNAseq data from d5-HSPCs.

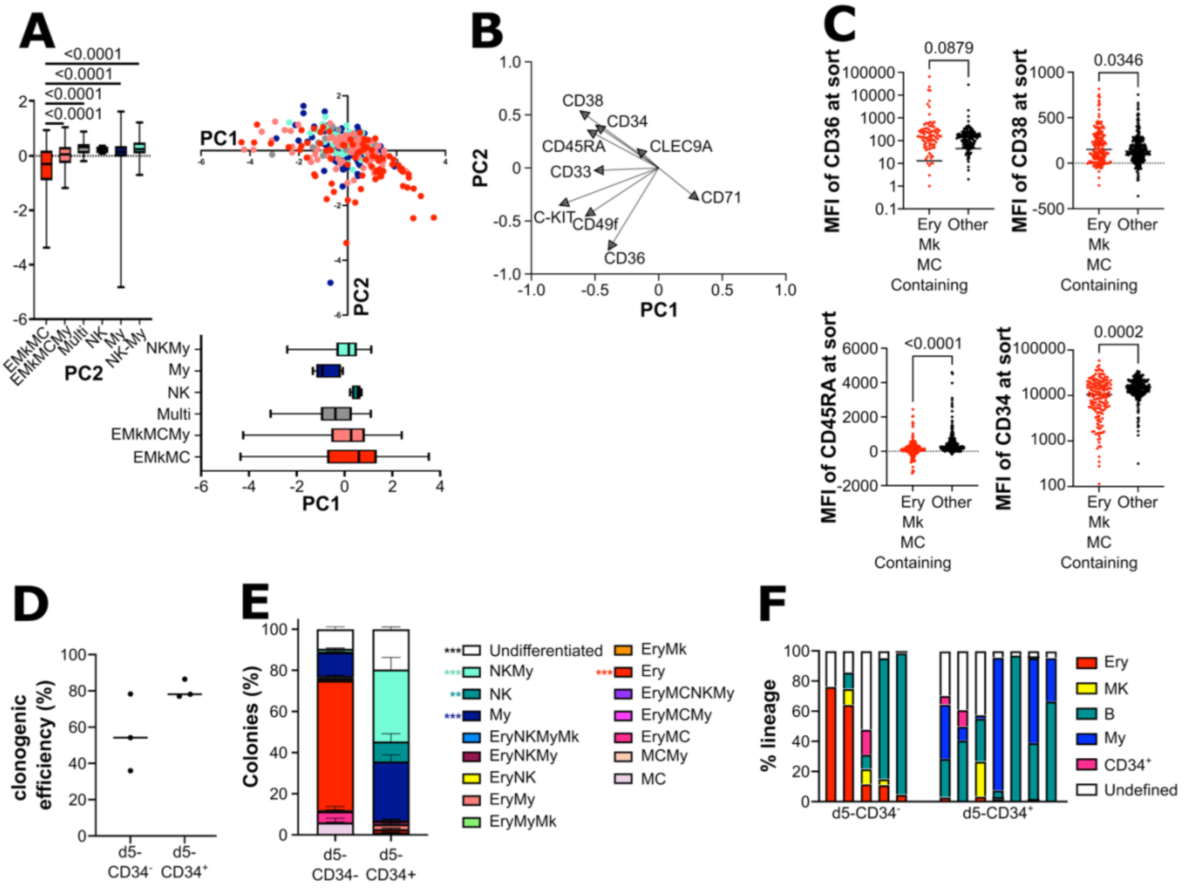


**Suppl. Figure 2: Prospective Purification and characterisation of cMEMPs. (A-C):** Analysis of colony type and cell surface marker at day 4 of 49f$^{+}$ HSC cultured for 5 days in the EPD assay then single cell sorted. (**A**) Principal component analysis (PCA) of the surface marker expression at the time of sort, Colours indicate the type of differentiated colony produced by each single-cell after culture, boxplots of PC1 and PC2 values from single cells producing indicated types of differentiated colonies; p=0.0775 for PC1 and p<0.0001 for PC2 by 1–way ANOVA with Tukey's multiple comparisons (shown). (**B**) PC loading of cell surface markers. (**C**) Median fluorescent intensity (MFI) of CD36, CD38, CD45RA and CD34 cell surface expression in colonies containing Ery, Mk and/or MC vs all other colony types; p values by unpaired t-test (n = 174 Ery-Mk-MC containing colonies and n=247 other colonies from 3 independent CBs). **(D-E)**: Colonies derived from d5-CD34$^{-}$ and d5-CD34$^{+}$ that were single cell sorted into differentiation cultures. (**D**) Clonogenic efficiency and (**E**) percentage of colonies containing differentiated cells of the indicated lineages; n=3 independent CBs, n= 464 and 687 colonies in d5-CD34$^{-}$ and d5-CD34$^{+}$ respectively. Mean ± SEM is shown, $p<0.0001$ by 2-way ANOVA with Sidak's multiple comparisons (shown). **(F)** Distribution of differentiated cell types from each indicated lineage in the human graft of individual mice (injected femur, each bar represents one mouse) engrafted with either d5-$CD34^-$ or d5-$CD34^+$ at 2 weeks post-transplantation.

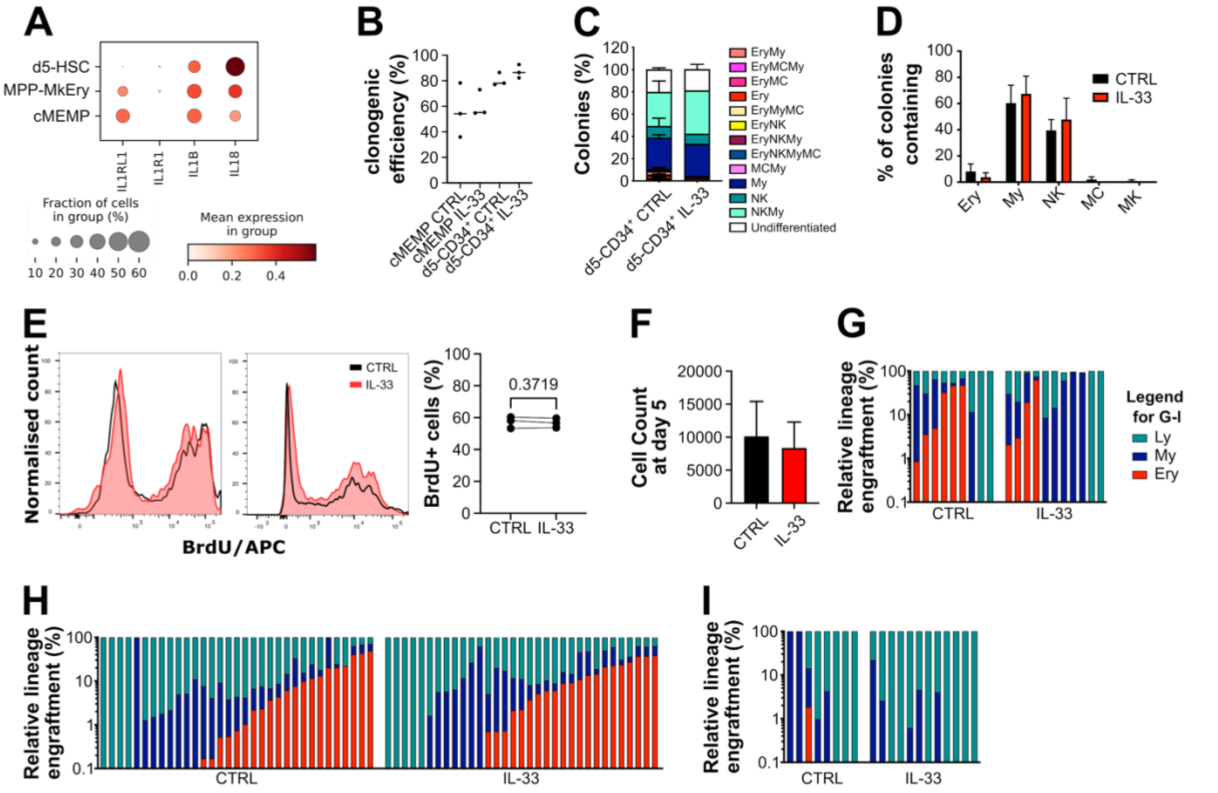


**Suppl. Figure 3: Functional effects of IL-33 in the EPD model. (A)** 10x genomics scRNAseq data from d5-HSPCs cultured in the EPD assay. Expression values of selected IL-1 family genes in the HSC, MPP$^{MkEry}$ and MEMP clusters as defined in Fig.1H. Circle colour shows mean expression values and circle size represents the proportion of cells expressing cells that gene per cluster. **(B-D)** 49f$^{+}$ HSCs were cultured in the EPD assay ±IL-33 then single cells from the indicated populations were sorted into the differentiation assay described in Fig. S1D. **(B)** Clonogenic efficiency, **(C)** percentage colonies of the indicated type and **(D)** percentage colonies containing specified lineages (**D**) n= 3 independent CBs. (**E**) Left panel: representative flow cytometry plots of BrdU staining of d5- HSPC obtained from EPD culture ± IL-33. Right panel: Percentage of Brdu$^{+}$ cells; n=3 p=0.3719 by paired-test. (**F**) Cell count of d5-HSPCs cultured in the EPD assay ± IL-33; n=7 p=0.3379 by paired t-test. (**G-I**): Distribution of differentiated cell types from each indicated lineage in the human graft of individual mice (injected femur, each bar represents one mouse) engrafted with d5-HSPCs derived ± IL-33 in the EPD assay at (**G**) 4 weeks and (**H**) 20 weeks after primary transplantation and after (**I**) 12-weeks post-secondary transplantation. My lineage: $CD45^{++}CD33^{+}$ cells; Ly lineage: $CD45^{++}CD19^{++}$ cells (positive for two distinct CD19 antibodies); Ery lineage: $GlyA^{+}$ cells.

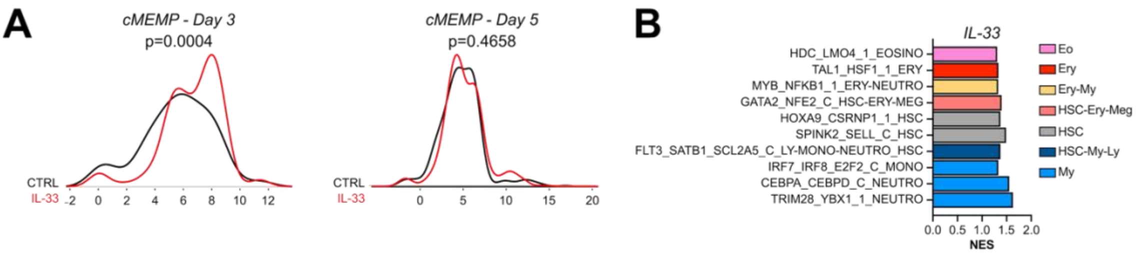


**C**

| Day | Group | Uni-, Bi- and Tri-Potent Primed | Non-Primed | Fisher's exact test two-tailed |
|---|---|---|---|---|
| 3 | CTRL | 94 | 54 | p<0.0001 |
| | IL-33 | 348 | 51 | |
| 5 | CTRL | 337 | 104 | p<0.0001 |
| | IL-33 | 502 | 32 | |

**D**

| Day | Group | Uni-Potent | Bi-/Tri-Potent | Fisher's exact test two-tailed |
|---|---|---|---|---|
| 3 | CTRL | 74 | 20 | p<0.0001 |
| | IL-33 | 196 | 152 | |
| 5 | CTRL | 197 | 140 | p<0.0001 |
| | IL-33 | 170 | 332 | |

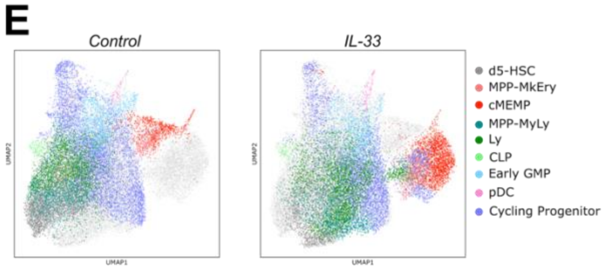


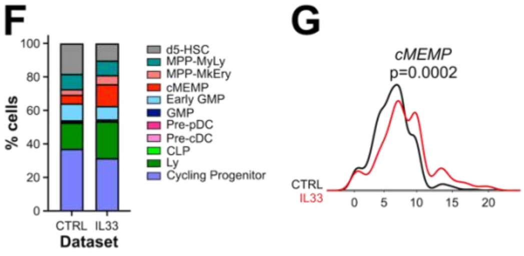


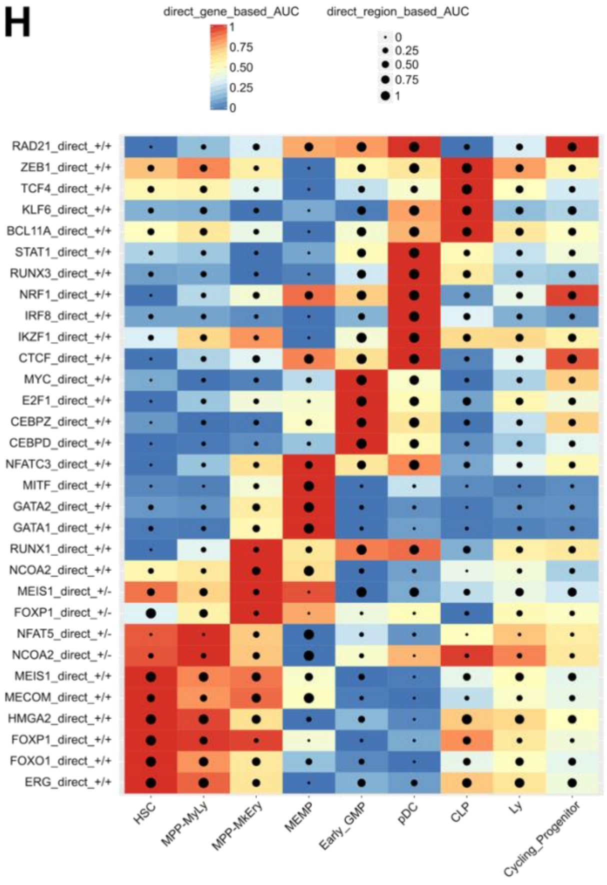


**Suppl. Figure 4: Effects of IL-33 on cMEMP transcriptional features. (A-D)** Analysis of 10x genomics scRNAseq data from 49f⁺ HSCs cultured ± IL-33 in the EPD assay for 3 or 5 days. n= 4921, 4357, 8402 and 3048 cells in day 3 control, day 3 IL-33, day 5 control and day 5 IL-33 respectively (after QC). (**A**) Distribution of cMEMP cells along the pseudotime ± IL-33 at day 3 (left panel) and day 5 (right panel). **(B)**

GSEA comparing day 5 cMEMP control vs IL-33, selected significant (FDR<0.05) lineage-priming gene sets from (Velten *et al.*, 2017) shown. Colour of the bars represents type of lineage. (**C-D**) Tables of counts of single cMEMP cells primed towards Ery, Mk or MC lineages (**C**), or predicted to be uni-, bi- or tripotent (**D**, see methods for assignment of lineage priming), p-values comparing control to IL-33 by Fisher's exact test. (**E-I)**: Analysis of 10x multiome (combined scRNA-seq and scATAC-seq) from 49f+ HSCs cultured ± IL-33 in the EPD assay for 5 days. n= 17,703 and 15,928 cells in control and IL-33 conditions respectively (after QC). (**E**) UMAP embedding based on RNA modality with annotated cell clusters shown. (**F**) Percentage of each cluster. (**G**) Distribution of cMEMP cells along the pseudotime ± IL-33 at day 5. (**H**) Heatmap-dotplot for a selection of eRegulons across the annotated clusters (CTRL and IL-33 conditions combined). The heatmap colour represents regulon activity, the size of the dotplot represents regulon accessibility. Statistics (panel **A** and **G**): Group comparisons were evaluated using a two-tailed permutation of medians (10,000 iterations) test. Reported p-values are unadjusted for multiple corrections.

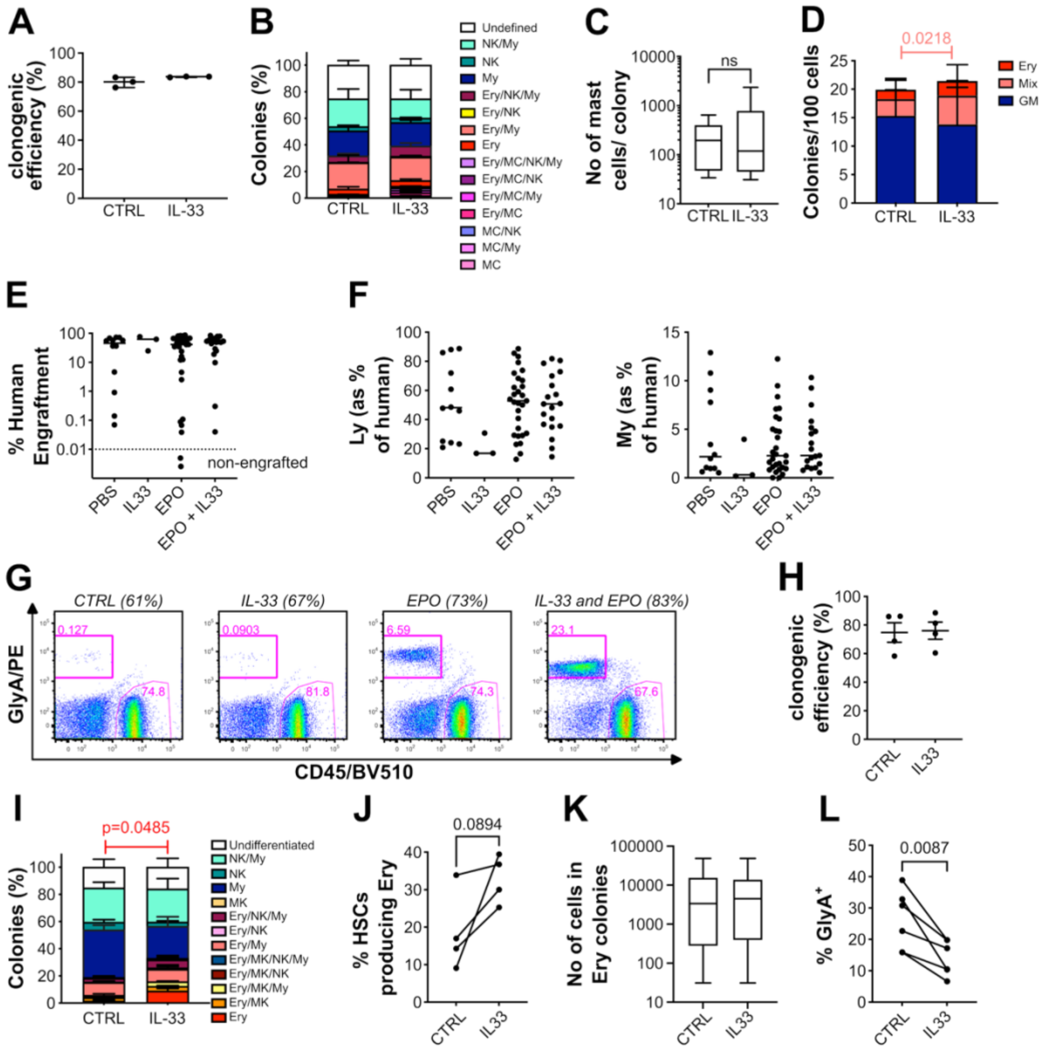


**Supp Fig 5: Effects of IL-33 downstream of cMEMP. (A-C):** Differentiation of single 49f$^+$ HSCs plated in MCEM medium. (**A**) Clonogenic efficiency. (**B**) Percentage of each colony type. (**C**) Number of mast cells per colony. n=3 independent CBs, n=537 control and n=562 IL-33 colonies for (**A-B**), n=14 control and 41 IL-33 colonies that produced MCs for (**C**). (**D**) Normalised number of colonies obtained from 49f$^+$ HSC plated directly into the CFU assay, n=12 independent CBs, statistics by paired t-test, mean ± SD shown. Colony types: Erythroid (Ery), granulocyte and monocyte (GM) or a combination of both (Mixed). (**E-G**): NSG mice engrafted with CB CD34$^+$ and then treated with PBS, IL-33, EPO or EPO and IL-33. (**E**) Percentage human engraftment, (**F**) Lymphoid (Ly, B cell) and Myeloid (My) engraftment as a percentage of human engraftment and (**G**) representative flow plots for GlyA and human CD45. n= 12, 3, 31 and 19 respectively, no statistical significance by one-way ANOVA. For (**E**) Dashed line: threshold of engraftment (%CD45$^{++}$ + %GlyA$^+$) ≥ 0.01 % and at least 30 cells recorded. Non-engrafted mice shown below dashed line. (**H-K)**: differentiation of single cell 49f$^+$ HSCs in MEM medium ±IL-33. (**H**) Clonogenic efficiency. (**I**) percentage of each colony type. (**J**) percentage single 49f$^+$ HSCs producing colonies containing Ery cells. (**K**) Number of Ery cells per colony. (**H-K**) n=4 independent CBs, n= 411 control and n= 445 IL-33 colonies. (**K)** n=84 control and 143 IL-33 colonies that produced Ery. (**I-J**) p-values by paired t-test. (**L**) Percentage GlyA$^+$ cells from PB Ery progenitors cultured for 8 days in medium as in (Swann *et al.*, 2020), n=6 independent PBs, p-vale by paired t-test.

**Suppl. Table 1: List of NSG mice used for xenotransplants** (excel file)

**Suppl. Table 2: Annotation and differential gene expression for Dataset 1** (excel file)

**Suppl. Table 3: Annotation and differential gene expression for Dataset 2 (d3/5-HSPC ± IL-33 RNA)** (excel file)

**Suppl. Table 4: Annotation and RSS scores for Dataset 3 (d5-HSPC ± IL-33 RNA+ATAC** (excel file)

**Suppl. Table 5: Antibody Panels for flow cytometry.** All antibodies from Biolegend or BD (marked with a *). All antibodies were titrated and validated using appropriate positive and negative control samples from human blood mononuclear cells.

| **Fluorochrome** | ***PANEL USE/* CELL SURFACE MARKER** | | | | | | | | | | | |
|---|---|---|---|---|---|---|---|---|---|---|---|---|
| | *HSC 1* | *HSC 2* | *d5-HSPC* | *d5-HSPC (index)* | HSC/MEP | *MEM* | *MCEM* | *Terminal Ery* | *Swann* | *Psaila* | *mouse BM 1* | *mouse BM 2* |
| **BV421** | CD7* | CD49f | CD33 | CD33 | | CD15* | CD15* | CD105 | | CD15* | | CD33 |
| **BV510** | Zombie (Aqua) | Zombie (Aqua) | Zombie (Aqua) | Zombie (Aqua) | Zombie (Aqua) | | FCER1a | CD41a* | | CD41a* | CD45 | CD45 |
| **BV650** | | | | C-KIT | | | C-KIT | | CD71 | | | C-KIT |
| **BV785** | | | CD19 | CD19 | CD123 | | | | | | | |
| **FITC** | CD45RA* | CD19 | CD71 | CD71 | CD41 | CD41 | CD41 | CD71 | CD71 | CD71 | CD19 | CD41 |
| **PE** | CD90 | CD45RA | CLEC9A | CLEC9A | CD45RA | GlyA* | GlyA* | GlyA* | GlyA* | GlyA* | GlyA* | GlyA* |
| **PECy5** | CD49f* | | | CD49f* | CD49f* | CD45 | CD45 | C-KIT | CD45 | CD45 | CD45 | CD45 |
| **PECy7** | CD38 | CD38 | | CD38 | CD38 | CD14 | CD14 | CD38 | | CD14 | CD14 | CD203c |
| **APC** | CD10 | CD90* | CD36 | CD36 | CD90* | CD56 | CD56 | CD36 | | CD42b | CD33* | CD61 |
| **Alexa 700** | CD19 | | CD45RA | CD45RA | CD19 | | | CD45RA | | | CD19 | CD19 |
| **APCCy7** | CD34 | CD34 | CD34 | CD34 | CD34 | CD11b | CD11b | CD34 | CD11b | CD11b | CD3 | CD34 |
